# Predicting the fMRI signal fluctuation with echo-state neural networks trained on vascular network dynamics

**DOI:** 10.1101/807966

**Authors:** Filip Sobczak, Yi He, Terrence J. Sejnowski, Xin Yu

## Abstract

Resting-state functional MRI (rs-fMRI) studies have revealed specific low-frequency hemodynamic signal fluctuations (<0.1 Hz) in the brain, which could be related to oscillations in neural activity through several mechanisms. Although the vascular origin of the fMRI signal is well established, the neural correlates of global rs-fMRI signal fluctuations are difficult to separate from other confounding sources. Recently, we reported that single-vessel fMRI slow oscillations are directly coupled to brain state changes. Here, we used an echo-state network (ESN) to predict the future temporal evolution of the rs-fMRI slow oscillatory feature from both rodent and human brains. rs-fMRI signals from individual blood vessels that were strongly correlated with neural calcium oscillations were used to train an ESN to predict brain state-specific rs-fMRI signal fluctuations. The ESN-based prediction model was also applied to recordings from the Human Connectome Project (HCP), which classified variance-independent brain states based on global fluctuations of rs-fMRI features. The ESN revealed brain states with global synchrony and decoupled internal correlations within the default-mode network.

## INTRODUCTION

Neural oscillations have been extensively studied in both animal and human brains from cellular to systems levels^1–4^. Power profiles of EEG signals, as well as slow cortical potentials (SCP), exhibit a slow oscillation feature (<1 Hz), which is related to brain states mediating memory, cognition and task-specific behaviors^5–7^. Resting-state functional MRI (rs-fMRI) studies have revealed low-frequency hemodynamic signal fluctuations (<0.1 Hz)^8–11^, which have been confirmed by intrinsic optical imaging^12^, laser-doppler-flowmetry^13^, and near-infrared spectroscopy^14^. In particular, specific spatial correlation patterns can be observed in the slow oscillation of the rs-fMRI signal, e.g. the default-mode network (DMN)^15–17^. Concurrent fMRI and electrophysiology studies have shown a correlation of the fMRI signal fluctuation with the EEG signal power profile and SCP low-frequency oscillations, which are candidates for neural correlates of the rs-fMRI signal^18–23^. In addition, the slow oscillation of rs-fMRI and hemodynamic signals from vessels are highly correlated to simultaneously acquired intracellular Ca^2+^ signal fluctuations in rodents^24–26^, which are higher-resolution correlates of the hemodynamic rs-fMRI signal.

Efforts have been made to interpret functional indications of the rs-fMRI spatial correlation patterns, including the dynamic correlation mapping scheme^27–29^, and arousal state-dependent global fMRI signal fluctuation studies^30–32^. Because of the high variability in different dynamic states, physiological and non-physiological confounding factors also contribute to the rs-fMRI low-frequency oscillation^33–35^. In particular, global fMRI signal fluctuations are one of the most controversial oscillatory features to be linked to dynamic brain signals^36–44^. For example, the rs-fMRI signal from the white-matter tract has been used as a nuisance regressor to remove the global noise contribution^45, 46^. Interestingly, simultaneous fMRI and EEG studies in the monkey brain demonstrate a strong linkage of brain state changes to the global rs-fMRI signal fluctuations. This phenomenon has been observed at the level of single-vessel fMRI dynamic mapping with concurrent calcium recordings, which show stronger neural correlation with the fMRI signal detected from individual penetrating vessels than the rest of voxels through the whole rodent cortex^24^. This highly coherent vessel-specific fMRI signal fluctuation is a direct signal source that is closely linked to global brain state changes. Here, we applied the artificial state-encoding neural network system in a prediction scheme to better model the brain state-specific coherent oscillatory features from the vessel voxels.

The echo state network^47^ (ESN), a recurrent neural network (RNN) based on reservoir computing^48, 49^, provides a computational framework for temporally predicting dynamic brain signals. The ESN’s main component is a dynamic reservoir consisting of recurrently connected computational nodes (neurons) that encode temporal patterns of input signals, i.e. the vessel-specific rs-fMRI signal, into a state matrix. The second element of an ESN is a linear decoder, which generates predictions based on reservoir’s internal states. ESNs have been successfully used for time series prediction^50^, estimating directed connectivity^51^ modeling nonlinear systems^52^ and superimposed oscillators^53^, and local cortical dynamics^54^. Other artificial neural networks have been applied to fMRI data to encode brain dynamics with the goal of characterizing psychiatric diseases^55, 56^, modeling task or sensory-evoked activation^57–59^ and decoding task or stimuli properties from fMRI activity^60, 61^. Artificial neural networks can depict dynamic brain signals over a range of time scales and contexts^62–66^.

In the present study, ESNs were trained to predict dynamic changes of single-vessel rs-fMRI signal fluctuations from the brains of rats and humans. fMRI recordings from single blood vessels^24, 67^ were used to extract highly correlated vessel-specific fMRI signals from venules or veins, which have been shown to be directly correlated to the underlying intracellular Ca^2+^ signal fluctuation from neurons in rat brains^24^. Vessel-specific fMRI signals were used as training data to extract highly correlated slow oscillation features with varied noise profiles and extract brain state-dependent global fMRI signals. The ESN reservoirs and decoders predicted the temporal evolution of slow oscillations of the fMRI signal 10 seconds into the future. The trained network differentiated the global fMRI signal fluctuation from the DMN-specific temporal dynamic patterns in the Human Connectome Project (HCP) data^68^.

## RESULTS

Two datasets were used in our study, one from rodents and another from humans. In the first part of the study, we trained an ESN to encode temporal dynamics of BOLD-fMRI signals from individual vessels in anesthetized rat brains to estimate the prediction efficiency. In the second part of the study, we trained an ESN to predict the slow oscillation of fMRI signals from occipital lobe sulcus veins of awake human subjects and applied the ESN trained on human data to classify the brain-state changes from rs-fMRI data acquired by the Human Connectome Project.

### Extracting slow oscillatory features of the single-vessel fMRI signal from rat brains

We used recordings obtained from a balanced steady-state free precession (bSSFP) sequence^69^ on single-vessel fMRI data from anesthetized rats^24^. Arteriole-venule (A-V) maps based on the multi-gradient-echo (MGE) sequence were acquired to localize individual venules penetrating the cortex, which were shown as dark dots due to the fast T2* decay of the deoxygenated blood (Fig. 1a)^67^. After registering functional data with the A-V map, fMRI time courses from individual venules were extracted and analyzed using independent component analysis (ICA)^70–72^. Fig. 1b shows the time series of the largest ICA component, which is dominated by the low frequency fluctuation (<0.1 Hz). The superposition of this ICA component with the single-vessel fMRI signal fluctuation on the A-V map overlapped with venule-dominated patterns (Fig. 1c). Fig. 1d shows the raw bSSFP-fMRI signal fluctuation from three venules, as well as their power spectral density (PSD) plots. These data presented highly coherent oscillatory features of single-vessel fMRI signals, which can be used as a training set.

**Fig. 1.**
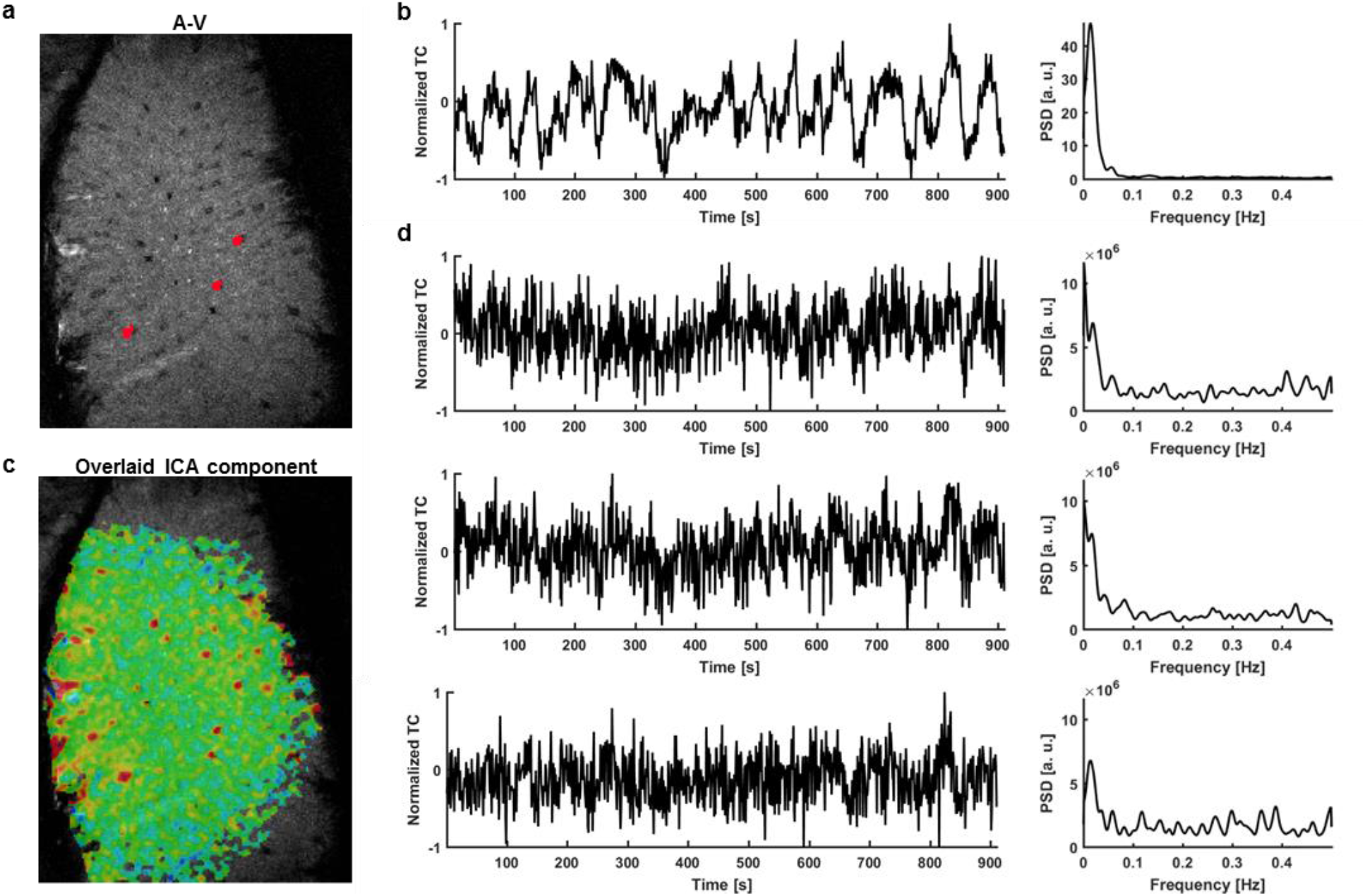
Extraction of signals from single venules exhibiting strong slow fluctuations – rat. **a**, The A-V map enables localization of single venules (dark dots) in the rat somatosensory cortex (red – 3 vessel masks; plotted in d.). **b**, Time course of the slowly changing ICA component shaping vascular dynamics and its power spectral density estimate (PSD). **c**, The corresponding ICA spatial map highlights the presence of slow fluctuations predominantly in veins. **d**, Examples of extracted vascular time courses selected for further processing (marked as red dots on the A-V map in *a*) along with their PSDs. The ICA component is present in the signals, but the noise level is much higher and individual differences are clearly visible.

### Supervised training of the ESN-based prediction of the fMRI slow oscillation

Fig. 2 illustrates the basic schematic of the ESN-based prediction. The single-vessel fMRI signals showing a strong slow oscillatory correlation (Fig. 1) were used as input time series for the supervised training. As described in the Methods section, a recurrent network-based reservoir was predefined to encode the state of the input signal’s temporal dynamics. As the key component of the reservoir, the state-weighting vector was optimized to produce output predictions based on supervised training. The targets of the output were bandpass-filtered fMRI signals from the voxels of the same vessel with a 10 s time shift. Pearson correlation analysis was performed to estimate the correlation coefficient (CC) between ESN’s output predictions and the filtered target signals, to measure of the ESN’s performance. We used random search^73^ and cross validation to find the set of hyperparameter values that produced the best performing echo state network (ESN) (Fig. S1, see details in the Methods section).

**Fig. 2.**
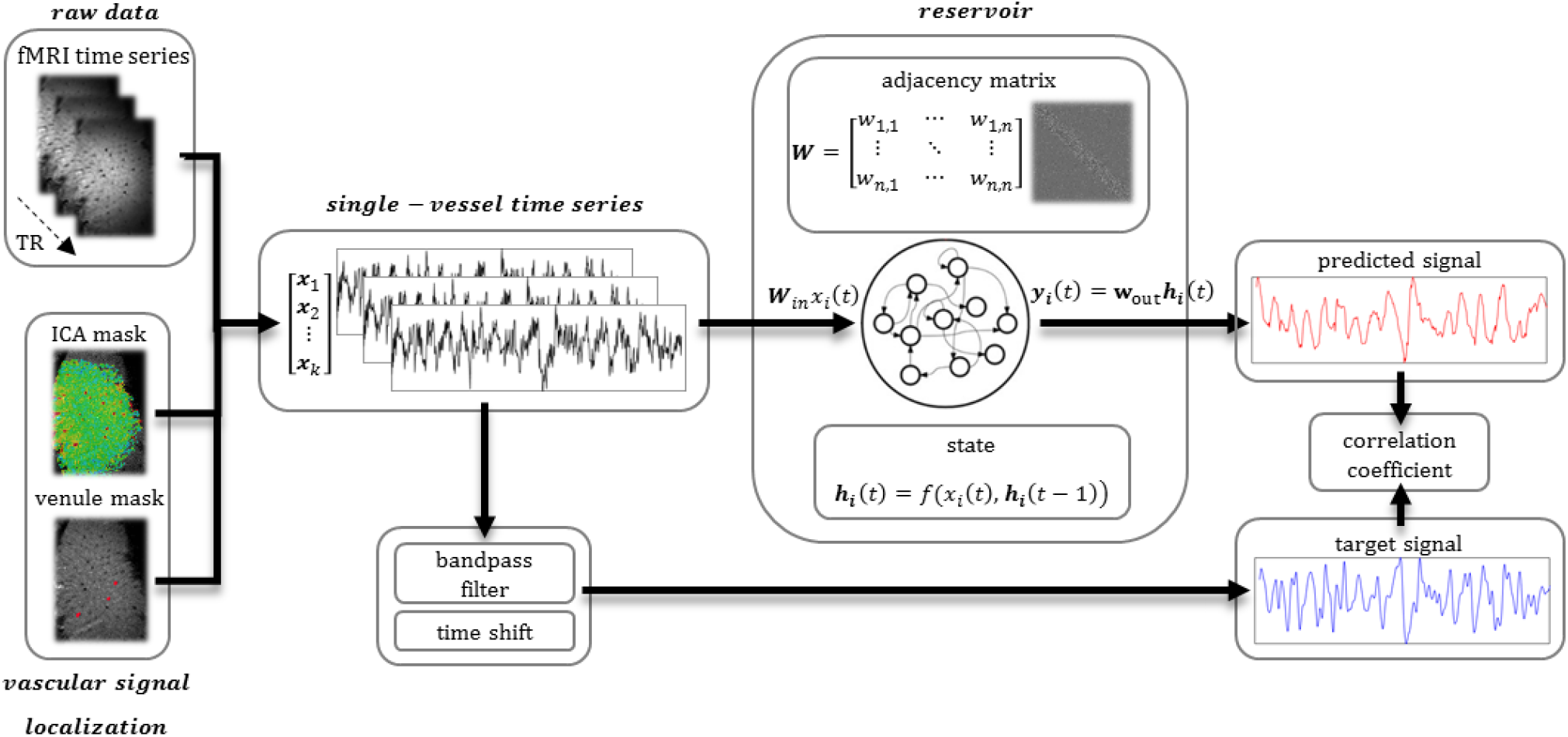
Prediction system operation pipeline. Raw vascular data are extracted from fMRI data using venule and ICA masks. These temporal signals are inputs to the ESN; they are also bandpass filtered and shifted by 10 seconds to become target outputs of the network. The reservoir encodes the temporal dynamics of input signals into state vectors. The decoder interprets these states and generates a prediction of the slow fluctuation’s value 10 seconds ahead. After generating the full predicted time series, the prediction is compared with the target output using Pearson’s correlation coefficient.

### ESN-based single-vessel fMRI slow oscillation prediction in anesthetized rats

We first illustrate the predictive capacity of the trained ESN by analyzing the correlation coefficients across all cross-validation tests. Fig. 3a demonstrates the CC of the slow oscillation prediction of all vessels from a representative rat. For each vessel, we generated a surrogate control time course that mimicked the frequency power profile of the fMRI signal. To differentiate the control dataset from true brain dynamic signals, we randomized the phase distribution of its frequency components^74, 75^ (Fig. S2, see Methods). The ESN prediction performance showed significantly higher mean CC for fMRI data (CC = 0.34 ± 0.01 s.e.m.) than surrogate controls (CC = 0.24 ± 0.01 s.e.m.) (Fig. 3b). Fig. 3c shows the predicted ESN time course from the vessel with the highest prediction score (CC = 0.53, t_lag_ = −2 s) in contrast to the surrogate control signal corresponding to the same vessel (CC = 0.32, t_lag_= −1 s). This shows that the trained ESN was better at predicting the fMRI signal fluctuations.

**Fig. 3.**
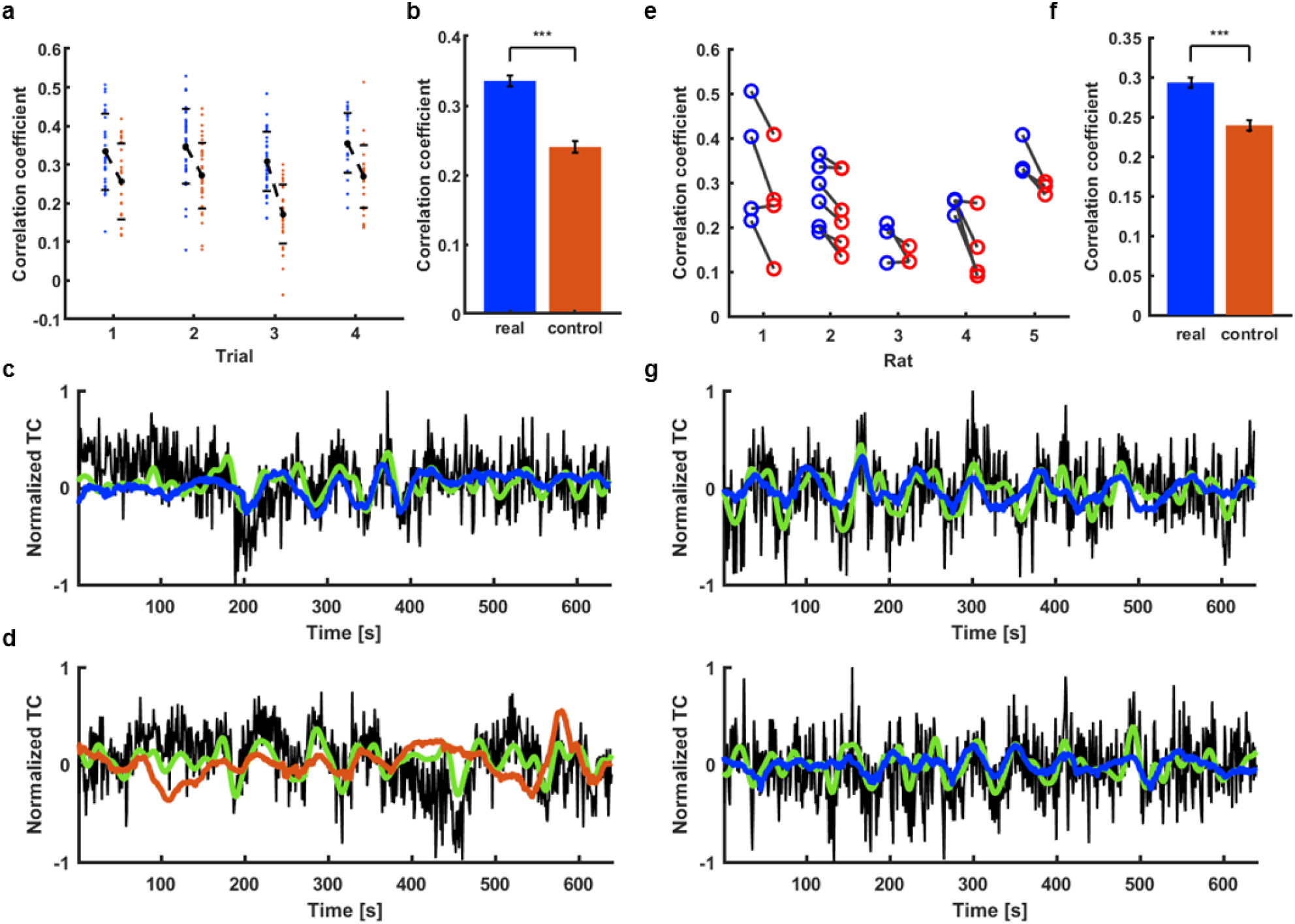
ESN prediction of the spontaneous slow fluctuation of rat vascular dynamics. **a**, Prediction scores of all the signals extracted from a single rat (blue dots) ordered by trials. Real data are matched with controls (red dots) for every vessel. Black dots show mean scores across trials and bars are SD values. **b**, Significantly higher mean of training rat real data prediction scores (CC=0.34 ± 0.01 s.e.m.) compared to controls (CC = 0.24 ± 0.01 s.e.m.; paired-sample t-test, p=1.7*10^-20^). **c**, The signal from a single vessel with the best prediction score (CC = 0.53, t_lag_=-2 s; black – raw data, green – target signal, blue – network prediction). **d**, Surrogate signal created to match the real vascular signal shown in c (CC = 0.32, t_lag_=-1 s; black – raw data, green – target, red – network prediction). **e**, Mean prediction scores for trials extracted from five rats (blue) and their corresponding controls (red). **f**, Significantly higher mean of different rats’ real data prediction scores (CC=0.29 ± 0.01 s.e.m.) than controls (CC = 0.24 ± 0.01 s.e.m.; paired-sample t-test, p=9.5*10^-26^). **g**, Predictions of single-vessel signals from two different rats (v1, CC = 0.55, t_lag_=-3 s; v2, CC = 0.55, t_lag_= 0 s).

In addition, the ESN trained on one rat was used to predict the fMRI fluctuation of five different rats. Fig. 3e demonstrates trial-specific plots of mean CCs from all vessels in comparison to their surrogate controls (380 vessels from 5 rats), showing significantly higher CC of the fMRI signal than that of surrogate controls (Fig. 3f). Fig. 3g shows predicted slow oscillatory time courses of two vessels from different rats based on the trained ESN (v1, CC= 0.55, t_lag_ = −3 s; v2, CC=0.55, t_lag_ = 0 s). These results indicate that the fMRI signal fluctuation can be predicted by the trained ESN.

### ESN-based single-vessel fMRI slow oscillation prediction in awake human subjects

As previously reported^24, 76^, the fMRI signal from sulcus veins of the occipital lobe demonstrated highly correlated slow-oscillatory features (Fig. 4). The vein-specific rs-fMRI signal fluctuations were recorded with high-resolution EPI-fMRI with 840 x 840 µm in-plane resolution and 1.5 mm thickness (Fig. 4a, veins are dark dots) and analyzed with ICA. The largest vascular ICA component exhibited slow oscillatory fluctuations in the 0.01 - 0.1 Hz frequency range (Fig. 4b) and its correlation map primarily highlighted the individual sulcus veins in the EPI image (Fig. 4c). Fig. 4d shows raw fMRI time courses from a few sulcus veins, demonstrating the vessel-specific time courses and PSDs with varied noise contributions to different veins. A difference in power distribution between species is visible in the PSDs. A significantly wider range of frequencies contribute strongly to time courses extracted from human vessels compared to rat data (Fig. S3, humans_FWHM_: 0.031±0.01Hz; rats_FWHM_: 0.008±0.001Hz, p=0.001). These results also enable the use of the ESN to encode the slow oscillation based on the vessel-specific fMRI signals from human brains.

**Fig. 4.**
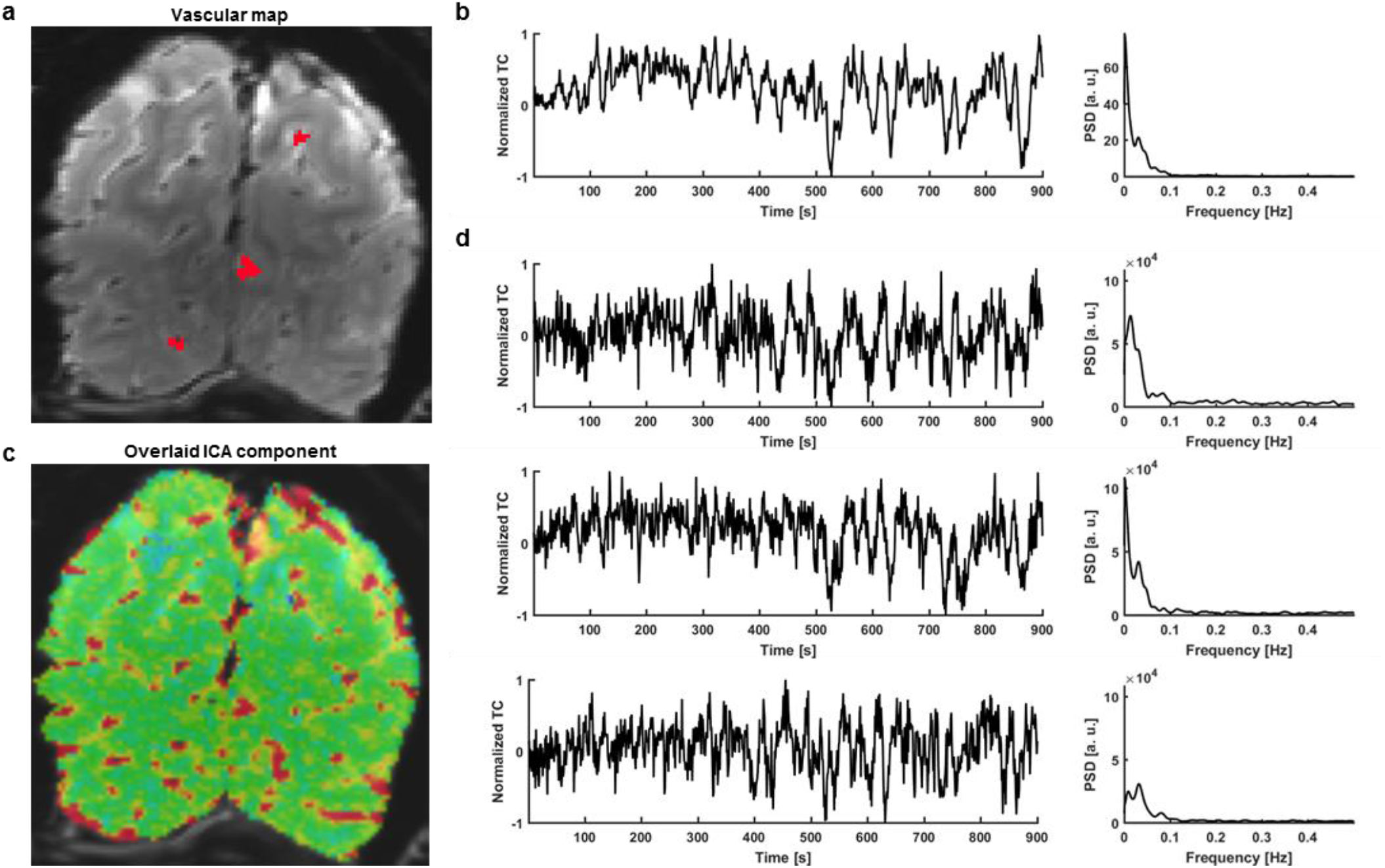
Extraction of signals from single veins exhibiting strong slow fluctuations in humans. **a**, The mean of a human single-vessel EPI time series enables the localization of single veins (black dots) in the occipital cortex (red – 3 vessel masks; plotted in d.). **b**, Time course of the slowly changing ICA component shaping vascular dynamics and its power spectral density estimate (PSD). **c**, An ICA spatial map highlights the presence of slow fluctuations predominantly in sulcus veins. **d**, Three single vessel time courses selected for further processing (marked as red dots in *a*) along with their PSDs. The ICA component is present in the signals, along with individual variations.

In contrast to the multi-trial single-vessel rat fMRI studies, only one trial (15 min) was acquired from each human subject (159 veins from 6 subjects). To perform the supervised training, we designed the 5+1 cross-subject validation process (trials from 5 subjects were used for training, and the sixth trial was used for test validation). Specific surrogate control time courses were created based on PSD profiles of fMRI signals acquired from individual veins in the human brain. Using the trained ESN, higher CC values were obtained by predicting slow oscillatory fMRI signals of individual veins compared to their surrogate controls (Fig. 5a), demonstrating a significantly higher mean CC value for brain dynamic signals (CC = 0.33 ± 0.01 s.e.m.) than for control datasets (CC = 0.27 ± 0.01 s.e.m.) (Fig. 5b). Also, the histogram of the cross-correlation lag times of the predicted and reference time courses showed the median of the lag time equal to 0, demonstrating the effective prediction. Figure 5d shows an example of a predicted slow oscillatory time course from a human subject based on the trained ESN (CC = 0.54, t_lag_ = −2 s). Fig. 5e shows the less accurate performance of the matching surrogate control (CC = 0.30, t_lag_ = 2 s). These results demonstrate the ESN-based cross-subject prediction of slow oscillatory fMRI signals.

**Fig. 5.**
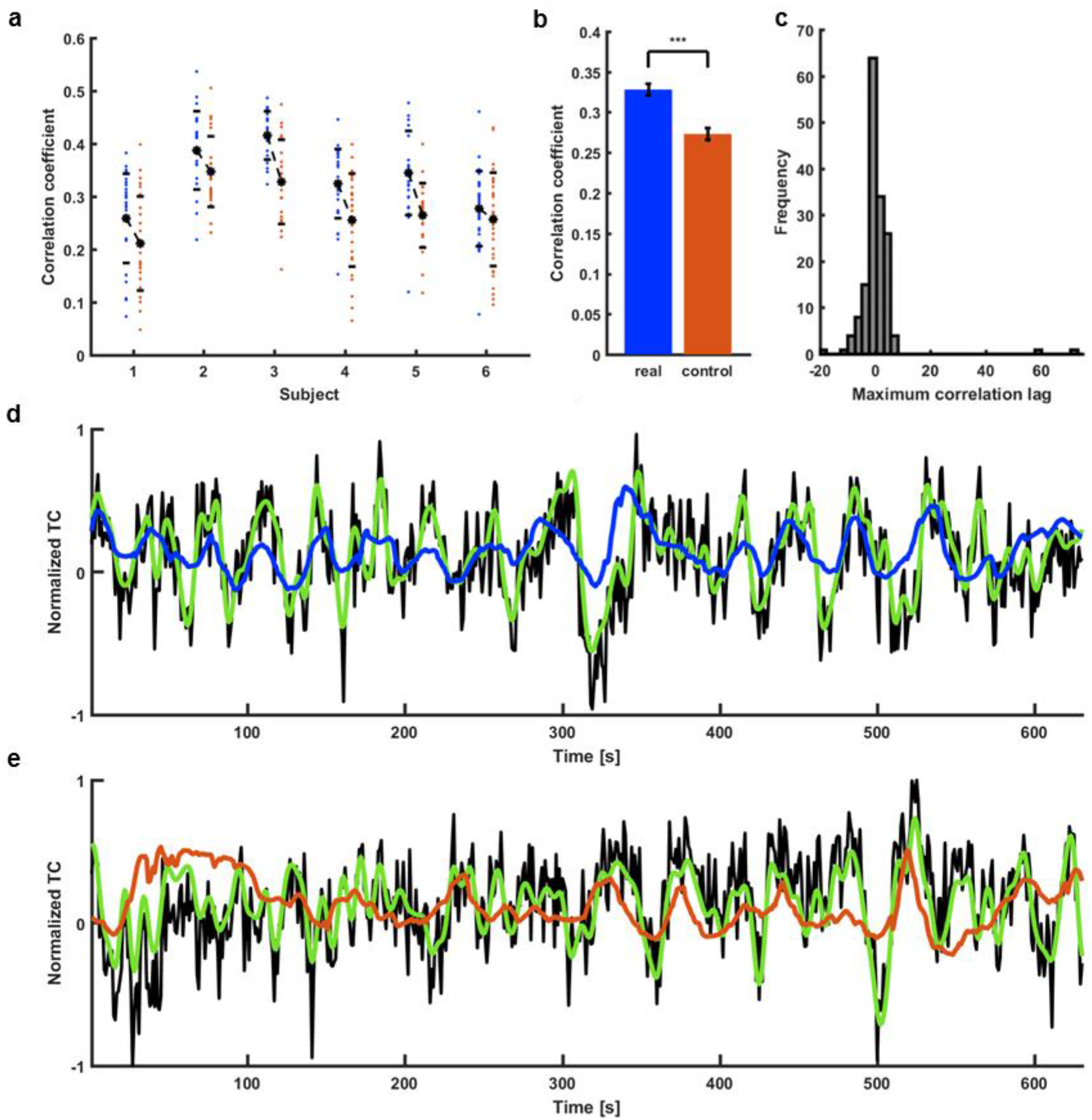
ESN prediction of the spontaneous slow fluctuation of human vascular dynamics. **a**, Prediction scores of all the signals extracted from 6 human subjects (blue dots). Real data are matched with controls for every subject (red dots). **b**, Significantly higher mean prediction score of real data (CC=0.33 ± 0.01 s.e.m.) as compared to controls (CC=0.27 ± 0.01 s.e.m.; paired-sample t-test, p=4.2*10^-14^). **c**, Histogram of lags at which the correlation between target outputs and network prediction was the highest. Distribution centered on 0 s (median = 0 s) indicates that the prediction wasn’t simply the filtered input. **d**, Prediction plot of the signal that obtained the highest score among all training human vessels (CC=0.54, t_lag_=-2 s; black – raw data, green – target, blue – network prediction). **e**, Prediction plot of the surrogate control signal created based on the real vascular signal shown in *d* (CC=0.30, t_lag_=-2 s; black – raw data, green – target prediction, red – network output).

The trained ESNs predicted artificial time courses with a range of peak frequencies and spectral widths (Fig. S4a,b). The predicted spread of the signal spectra preference for the ESN_human_ was greater than for ESN_rat_ as shown in the two-dimensional graphs of peak vs. width of the CC distribution (Fig. S4c,d). These species differences may reflect the difference in their rs-fMRI. Interestingly, the harmonic patterns had negative correlations for the preferred frequency, which could be a consequence of the trained ESNs favoring the dominating frequency ranges with the 10 s prediction interval.

### ESN-based prediction of the fMRI slow oscillation in the visual cortex (V1) of HCP data

Previously, we showed that smoothed single-vessel rs-fMRI correlation maps mimic conventional correlation maps in the human occipital area^24^. As shown in the PSD plots (Fig. 4), the vessel-specific fMRI slow oscillation dominates the 0.01-0.1 Hz frequency range. To examine whether the ESN trained by the single-vessel fMRI scheme can be used to predict the fMRI slow oscillation of a broader range of datasets, we applied the trained ESN to predict the rs-fMRI signals from the V1 of HCP data (a total of 3,279 rs-fMRI sessions; V1 signal extracted from left and right hemispheres separately, yielding 6,558 time courses resampled at 1 s TR, details in the Methods section). To examine the predictive capacity of the ESN on each trial of the HCP dataset, the CC of all prediction trials were plotted in a histogram. The CC distribution resembled a normal distribution centered on 0.28 (median) (Fig. 6a).

**Fig. 6.**
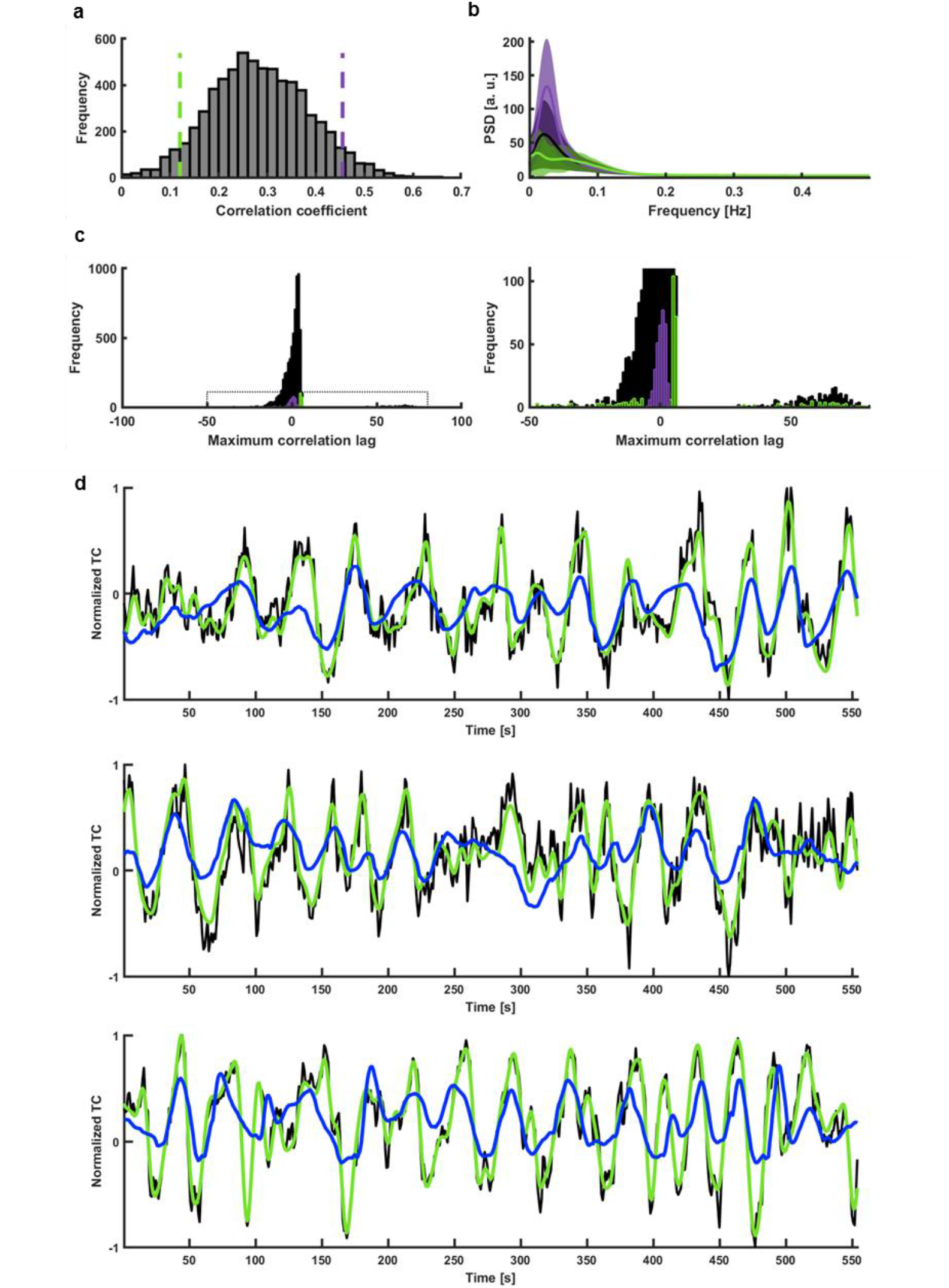
ESN categorization of V1 temporal patterns. **a**, Histogram of prediction scores obtained by predicting slow fluctuations of 6558 single-hemisphere V1 ROI signals extracted from HCP data. The used ESN was trained on occipital cortex single-vessel signals of 6 in-house subjects. Green and violet dashed lines mark the bottom and top 5% of correlation coefficients. **b**, Mean PSDs of time courses whose predictions obtained the bottom 5% (green) and top 5% (violet) scores. Shaded areas show s.d. **c**, Histogram of lags at which the correlation between targets and network outputs was the highest. The spread of values and a high number of large lags indicates a poor overall quality of prediction. However, the lags of top 5% of the predictions (violet) are concentrated around 0. The lags of bottom 5% (green) are spread across the highest and lowest lag values. *Right:* Enlarged region marked on the left plot. **d**, Predictions of signals with the three best correlations (CC_1_ = 0.65, t_lag,1_ = −1 s; CC_2_ = 0.61, t_lag, 2_= 2 s; CC_3_ = 0.61, t_lag,3_=2 s; black – raw data, green – target, blue – network prediction).

Next, we selected two clusters of the HCP dataset based on their CC (top 5%, bottom 5%), showing the top 5% trials with high power levels and the bottom 5% trials with low power levels at the 0.01 - 0.1 Hz frequency range (Fig. 6b). The lag time distribution of the top 5% group is centered at zero, unlike the bottom 5% group covering the whole range of lag values (Fig. 6c). In particular, many lag values of the poorly predicted sessions show a delay of more than the full wavelength of ESN’s preferred frequency. Fig. 6d shows three predicted slow oscillatory time courses from the HCP rs-fMRI sessions (top 5% group) (CC_1_ = 0.54, t_lag,1_= −2; CC_2_ = 0.61, t_lag,2_ =2; CC_3_ = 0.61, t_lag, 3_= 2). The predictions of the ESN were dominated by the low-frequency power in the rs-fMRI signals from individual trials.

### ESN-based brain state classification from HCP data

Here, we analyzed whole-brain correlation patterns of the HCP dataset, mainly focusing on the top and bottom 5% datasets from ESN predictions. Fig. 7 shows flattened cortical difference maps of seed-based correlations calculated for the two groups of HCP datasets. First, rs-fMRI time courses from the V1 ROI, the whole cortex (global mean), and the DMN were used to calculate voxel-wise correlation maps for all trials in the top and bottom 5% groups. These were then group-averaged and subtracted to create the correlation difference maps (Fig. 7a-f, the representative time courses from 4 subjects in each group were shown in Fig. S5). Similar differential patterns were detected for V1 ROI and global mean time courses, showing significantly stronger correlation pattern for the top 5% group covering most of the cortical regions (Fig. 7a-d). Although there is higher correlation to other cortical regions in the maps with the DMN-specific seed, no significant correlation differences were detected among DMN areas between the top and bottom 5% groups (Fig. 7e, f).

**Fig. 7.**
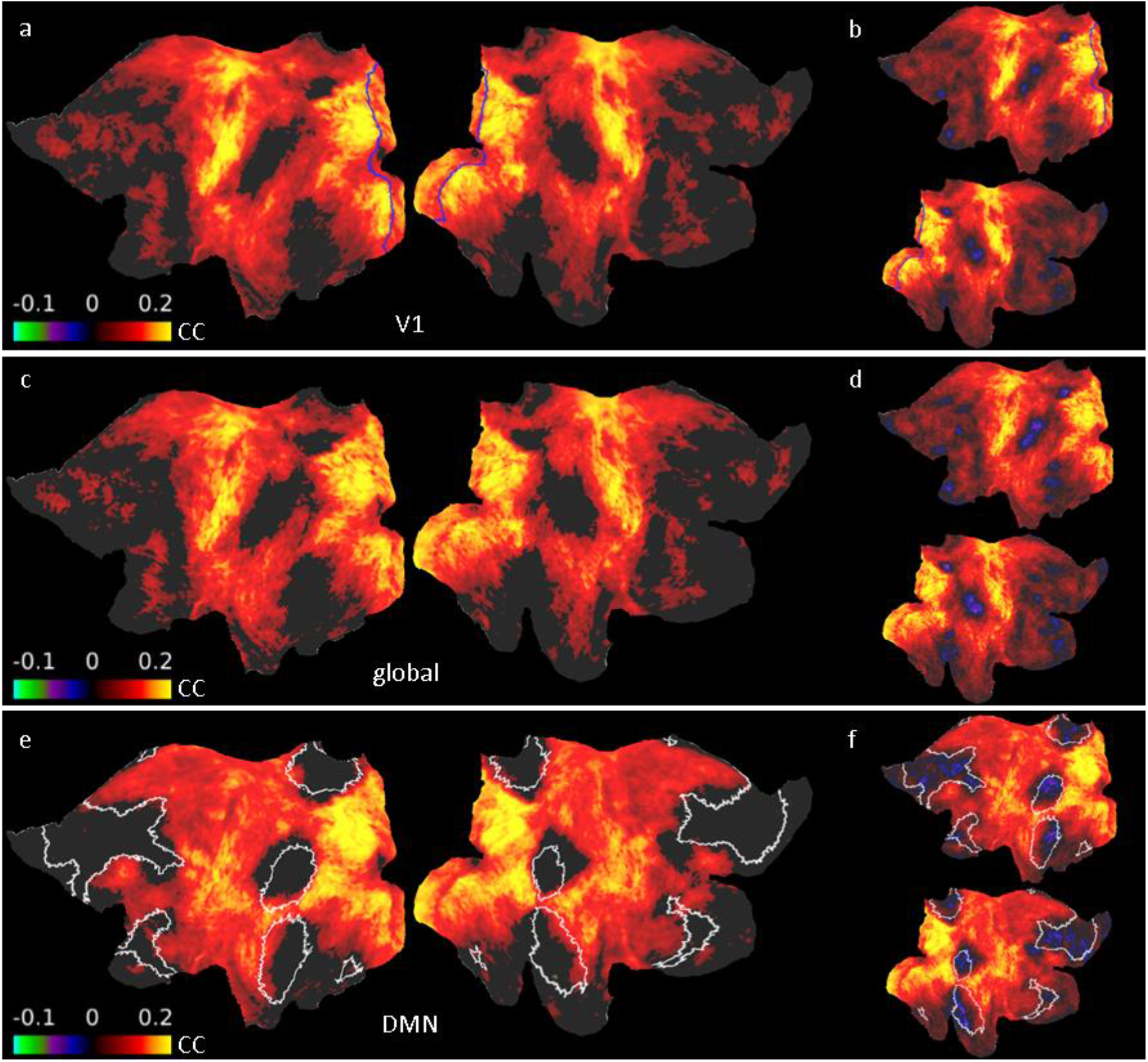
Difference maps for ROI seed-based correlations between well and poorly predicted fMRI sessions. **a**, The V1 seed region is marked with a blue border. Visual, sensorimotor and auditory areas display high increases in correlation. Nodes in which the difference was insignificant are masked. **b**, Same as a but without the mask. **c**, The average time course of the whole cortex served as the seed. The map resembles the result generated using V1 seeds (shown in a), suggesting that V1 signals extracted from the “top” group were mostly driven by the global signal. Nodes in which the differences were insignificant are masked. **d**, Same as c but without the mask. **e**, The average time course of the DMN ROIs served as the seed (marked with white borders). Nodes in which the difference was insignificant are masked. Despite showing increased synchrony with the areas dominated by the global signal, ROIs constituting the DMN don’t show significant differences between the groups. **f**, Same as e but without the mask.

We used DMN and visual ICA component time courses to compare their seed-based difference maps in order to better characterize the brain-state specific differences among resting-state networks classified by ESN predictions. The visual ICA-based map resembled the V1 ROI seed-based result (Fig. 8a-c). The DMN ICA signals are free of global signal contributions and represent intrinsic DMN activity. Using them enables the characterization of DMN-specific correlation patterns independent of the increase in global synchrony. The DMN ICA-based correlation maps of the top 5% subjects had reduced correlation, in particular showing significantly lower correlation features inside the DMN nodes compared to the bottom 5% sessions (Fig. 8d-f). The distinction of DMN-specific inter-network correlation was also presented in the correlation matrices based on the 360 ROIs predefined from the brain atlas^77^ (Fig. S6a). Correlation matrices computed for subcortical ROIs show increased correlation between the hippocampus and the brainstem with the global signal (Fig. S6b). These results indicate that the ESN-based classification can be used to differentiate the brain-state dependent rs-fMRI signal fluctuations in the HCP datasets.

**Fig. 8.**
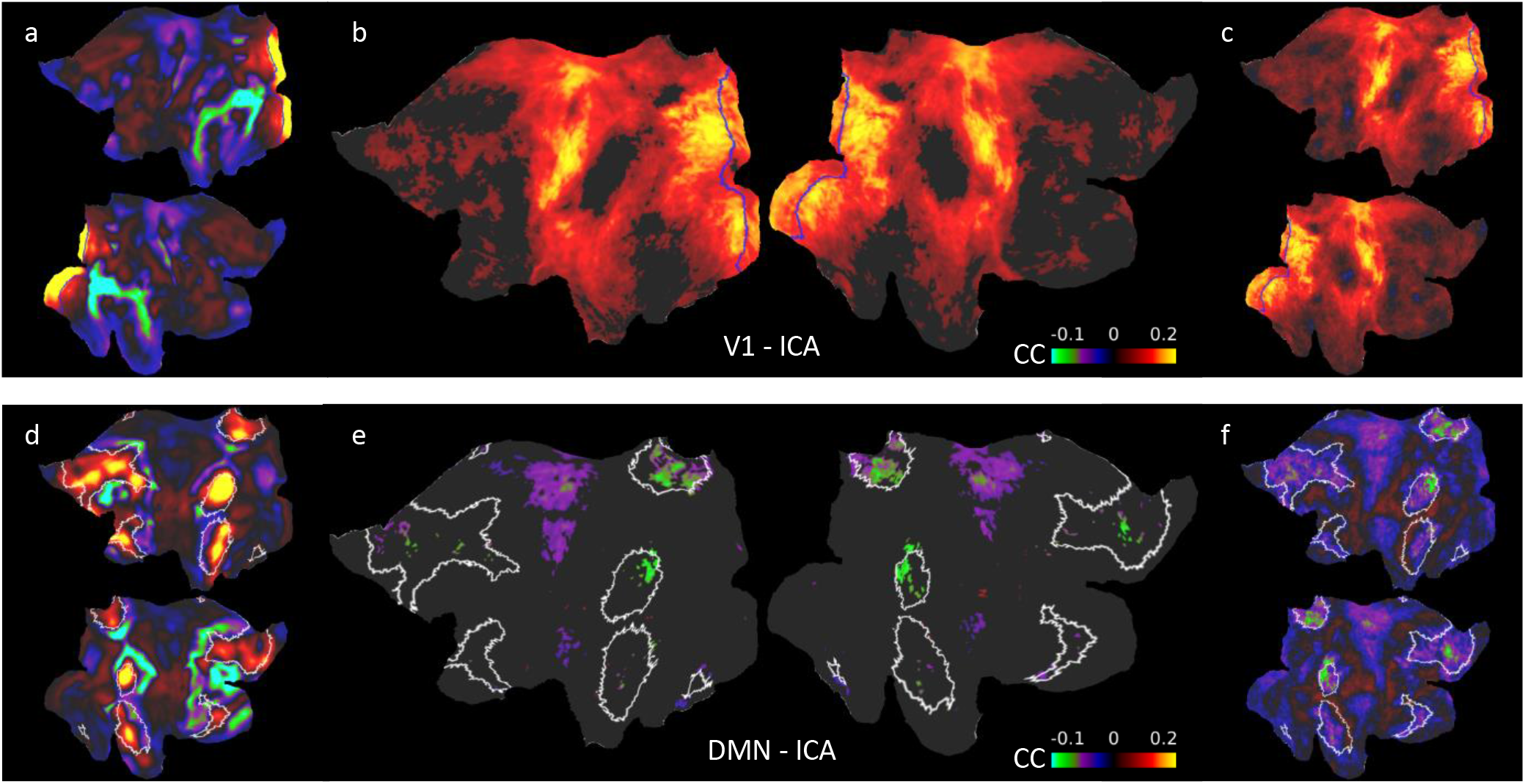
Difference maps for ICA seed-based correlation between well and poorly predicted fMRI sessions. **a**, V1 ICA component spatial map. V1 ROIs are marked by blue borders. **b**, Flattened cortical map showing the difference between the mean seed-based correlation maps of the “top” and “bottom” groups. The time course of the ICA component shown in a served as the seed. V1 ROIs are marked by blue borders. The connectivity pattern of the V1 ICA component resembles the pattern obtained by using the V1 ROI as the seed. Nodes in which the difference was insignificant are masked. **c**, Same as b but without the mask. **d**, DMN ICA component spatial map. DMN ROIs are marked by white borders. **e**, Flattened cortical map showing the difference between the mean seed-based correlation maps of the “top” and “bottom” groups. The time course of the ICA component shown in c served as the seed. DMN ROIs are marked by white borders. Nodes in which the differences were insignificant are masked. The intrinsic DMN signals show significantly reduced connectivity with DMN areas. **f**, Same as e but without the mask.

The classification scheme is not simply based on the variance of the rs-fMRI signal fluctuation. In contrast to the CC_ESN_-based classification (top vs. bottom 5%) of the HCP datasets, we also identified two groups of sessions with top vs. bottom 5% V1 variance in the same dataset (Fig. S7a-c). The CC values of the rs-fMRI variance-dependent groups for ESN-based predictions cover the total distribution range. In particular, the top and bottom 5% variance groups had much broader CC_ESN_ values and largely overlapped each other in the histogram plot (Fig. S7a). Similarly, the variance values of trials with the top and bottom 5% of ESN-predicted CC scores overlap and cover the whole range of the variance distribution (Fig. S7c). Also, no significant reduction of the internal DMN correlations were observed between the two variance-based groups (Fig. S7d-f). These results indicate that the ESN trained with vessel-specific rs-fMRI signals encodes specific brain state dynamic changes, which is less dependent on the variance of the rs-fMRI signal fluctuation.

In addition, we compared the ESN-based predictions with two other prediction schemes, the autoregressive moving average with exogenous input (ARMAX) modeling^78, 79^ and RNNs trained with the backpropagation algorithm^80, 81^. The best ARMAX models were found using an exhaustive grid search. The architectures and hyperparameters of the backpropagation-RNNs, the gated recurrent unit (GRU)^82^ and the long short-term memory (LSTM)^83, 84^ networks, were obtained using Bayesian optimization^85, 86^ (see Methods). The ESN obtained better prediction scores than ARMAX and showed very similar prediction performance to the other RNNs on our in-house datasets (Human: CC_ESN_ = 0.328 ± 0.01, CC_GRU_ = 0.324 ± 0.01, CC_ARMAX_ = 0.299 ± 0.01; Rat: CC_ESN_ = 0.304 ± 0.01, CC_LSTM_ = 0.305 ± 0.01, CC_ARMAX_ = 0.263 ± 0.01; mean ± s.e.m.) (Fig. S8a), as well as on the HCP datasets (Fig. S8b). We further compared the classification performance on the HCP datasets among the three different methods. The ESN and the backpropagation-RNN (i.e. the GRU) presented consistent group classification outcomes, showing similar distributions of the correlation coefficients of the top and bottom 5%, which was not the case with the ARMAX method (Fig. S8c,d). Also, the DMN ICA-based correlation differential maps from ESN and GRU methods showed a similar spatial pattern with significantly reduced correlation inside the DMN nodes of top 5% in comparison to the bottom 5% sessions, which was not detected by the ARMAX method (Fig. S8e). These results further confirmed the reliability of the RNN methods (both ESN and the backpropagation based RNNs) to classify the brain state-specific rs-fMRI signal fluctuations.

## DISCUSSION

We used the time courses of single-vessel rs-fMRI signals as inputs to train ESN networks to predict the rs-fMRI signal 10 s ahead in both rodents and humans. We also showed that the single-vessel fMRI-based training leads to ESN encoding specific to global fMRI signal fluctuations. The trained network was used to analyze HCP datasets with diverse brain states. For example, it allowed us to identify sessions with strong global synchrony and to decouple the global signal fluctuations from internal DMN correlations.

We selected the input fMRI time series from individual vessel voxels based on a previously established single-vessel fMRI mapping method^24, 67^. The BOLD fMRI signal has a direct vascular origin based on the oxy/deoxy-hemoglobin ratio changes^87–89^. The high-resolution single-vessel mapping method allows us to directly extract the venule-dominated BOLD signals with a much higher contrast-to-noise ratio (CNR) than the conventional EPI-fMRI integrating the BOLD signal from both tissue and vessels in large voxels^24, 67, 90, 91^. Although different vessel voxels may present cardiorespiratory noises, e.g. the respiratory volume change^33, 92^ or the heartbeat variability^93, 94^, a recent simultaneous fMRI and fiber-optic calcium recording study showed strong correlation of the major ICA vascular component of the rs-fMRI signal fluctuation (Fig. 1) with the calcium signal oscillation^24^. Also, these global hemodynamic signal changes are directly correlated with the calcium signal fluctuation through the whole cortex based on optical imaging^25, 26^. Thus, the global fMRI signal fluctuation detected from individual vessels represents changing brain states, and not the non-physiological confounding artifacts uniformly distributed through the brain, e.g. the respiration-induced B0 offset^95^ or other sources^34, 96^. In comparison to the voxel-wise or ROI-based time courses from low-resolution EPI images, the single-vessel rs-fMRI signal provides highly selective datasets for the supervised ESN training to encode brain-state dependent global fMRI signal fluctuations.

The predictions from the trained ESN’s vary across vessels as well as across trials. To validate this measurement, we used surrogate controls designed using the IAAFT method^75^. For every vessel, we generated an artificial signal showing a similar frequency power profile (Fig. S2) to its corresponding single-vessel rs-fMRI time course, but with randomized phases of complex Fourier components. It has been shown that high-frequency EEG power profiles are highly correlated to the low-frequency EEG signal fluctuation, i.e. phase-amplitude coupling (PAC), in both cortical and subcortical regions for a variety of brain states^97–102^. This feature has also been used for the correlation analysis of the concurrent EEG and rs-fMRI signal recordings from animals and humans^18, 19, 21, 23, 103–105^. Our analysis confirms that the phases of the slow oscillatory rs-fMRI signal carry critical dynamic brain state features^3^. By randomizing the phases, the surrogate control excludes dynamic brain features but preserves a high similarity in terms of the signal amplitude/power spectral distribution and autocorrelation structure for the verification of the ESN encoding. Also, the spectral characteristics of the ESNs demonstrate different preference maps in terms of the center frequency and the bandwidth depending on the training data from either rat or human data (Fig. S4). These training data showed differences in frequency power profiles given the inter-species diversity^106^ and the presence of anesthetics^24–26, 107–109^.

The global rs-fMRI signal is a critical confound of correlation analysis with many contributing factors from both physiological and non-physiological sources. In particular, whether the global mean fMRI signal should be removed before the analysis, which can create spurious correlation features, has been debated ^36–42, 44^. Also, the global rs-fMRI signal can over-shadow specific intrinsic RSN features, e.g. the anti-correlation of the DMN and task-positive RSNs^110–112^. One intriguing observation based on ESN-predicted results shows that the internal DMN correlations of the top 5% ESN performance group are reduced compared to the bottom 5% group (Fig. 8d-f), which is opposite to changes in the global correlation through the whole brain (Fig. 7, 8). It is noteworthy that the decreased internal DMN correlations are not visible through variance-based approaches (Fig. S7d-f). Thus, independent of the variance analysis method, the ESN-based approach reveals brain-state specific rs-fMRI signal fluctuations in the HCP datasets.

The contrast between internal DMN correlations and whole brain correlation patterns supports other sources of evidence that the global signals are dissociated from intrinsic brain network correlations^32^. Turchi et al. showed that the global rs-fMRI signal fluctuation can be directly modulated by inhibiting the activity of the basal forebrain nuclei, indicating that arousal leads to global rs-fMRI signals^32^. Global rs-fMRI signal fluctuations are also correlated with whether the eyes are open or closed^113–116^ and pupil dilation^117, 118^, and dynamic brain state changes that occur during different sleep stages^11, 63, 119–122^. A recent concurrent fMRI and calcium recording study has shown that the rs-fMRI fluctuation can be regulated by the arousal ascending pathway through the central thalamic nuclei and midbrain reticular formation^31^, implicating the subcortical regulation of the rs-fMRI signal fluctuation as previously reported from both non-human primate and human rs-fMRI studies^30, 32, 43^. Importantly, we also observed that the single-vessel rs-fMRI signal is specifically coupled to the global neuronal signal fluctuation^24^, which supports our single-vessel ESN training scheme to encode the brain-state specific global rs-fMRI signal fluctuations.

Thus, the ESN-based approach provides a variance-independent scheme to differentiate the global rs-fMRI fluctuation of the dynamic brain states. A promising direction for future work involves applying the proposed method to study the predictability of slow fluctuations in brain regions other than sensory cortices and to investigate which factors, besides arousal-related brain state changes, drive the predictions. Extending the platform to process whole-brain signals would provide a more synoptic view of the regularities present in brain dynamics in different states. Finally, the method could be integrated into a real-time fMRI platform to provide feedback stimuli and close the loop.

## METHODS

### Echo state network

To encode the dynamics of spontaneous slow fluctuations we used the echo state network (ESN)^47^, a recurrent artificial neural network belonging to the class of reservoir computing methods^48, 49^. It is trained by supervision. Its two main components are a dynamical reservoir encoding temporal patterns of input signals and a linear readout which decodes the reservoir’s state to generate the network’s output. A toolbox (http://minds.jacobs-university.de/research/esnresearch) providing basic ESN functionality was used and further developed for the sake of this work.

### ESN – state

The reservoir is a network of recurrently connected computational units called neurons. A state value is associated with each neuron. The states of reservoir neurons are driven by an external input as well as previous state values. This is illustrated by the equation:

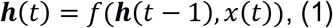

where ***h***(*t*) is the vector of reservoir states, *x*(*t*) is the input to the network and *f* is the activation function of each model neuron. This formulation leads to the network having a memory capacity spanning over multiple past inputs and allowing the reservoir to be described as a temporal kernel projecting the input time series into a high-dimensional feature space, whose dimensionality is equal to the number of reservoir’s neurons. Data transformed into this space are then decoded by the readout component of the ESN, which generates output predictions.

The reservoir is characterized by its weighting matrices ***W***, ***W***_***in***_, ***W***_***fb***_ and by the activation function *f*. Here, *f* was the hyperbolic tangent function. The matrices specify internal connections between reservoir elements (***W***), connections between the inputs and the reservoir (***W***_***in***_) and feedback connections from the readout into the reservoir (***W***_***fb***_). These parameters allow to formulate the basic update rule of ESN’s state:

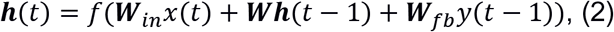

where ***y*** is the output of the network. The state update equation was further extended by incorporating leaky integrator neurons^123^ to enhance the network’s memory capacity. In this case, an additional parameter called the leaking rate *a* controls the fraction of the state which is preserved in the subsequent state and *γ* is a gain parameter brought about by discretizing continuous-time dynamics of leaky-integrator ESN equations:

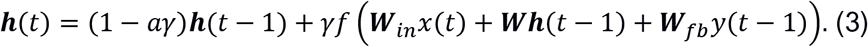

It is important to note that the input *x* might be extended by a constant input bias so that besides the vascular input, every neuron is also driven by a constant at every time step.

### ESN – readout

To generate the prediction, the decoder processes the ESN’s state. In the case of employing a linear readout, the output of the network is computed as:

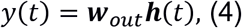

where ***w***_*out*_ is the output vector weighting the ESN’s state, which is learned by supervision. Once the reservoir has been fixed, input data are fed in it to generate the states matrix ***H*** according to eq. (4). The output vector is then computed using the target outputs ***Y***_*tr*_ and the matrix ***H*** containing reservoir states generated using all the training inputs:

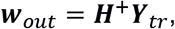

where ***H***^+^is the Moore-Penrose pseudoinverse of the states matrix ***H***.

### Random search optimizati

For a given ESN, the weight matrices ***W***, ***W***_***in***_, ***W***_***fb***_ defining its reservoir remain fixed after being initialized. They are not changed through training. To initialize the reservoir, a few hyperparameters need to be specified^124^. The performance of a single ESN instance is largely dependent on the choice of these hyperparameters and the randomness involved in weight initialization. There is no set of parameter values that would lead to good ESN performance on all problems posed. As the process of training a single ESN is not computationally expensive compared to other methods like e.g. network training using backpropagation, many reservoirs might be evaluated. Random search is a simple yet effective way of exploring the hyperparameter space for a given task^73^. For every parameter whose value isn’t fixed, a range of possible values is specified and every time an ESN is generated, the parameter values are selected at random from the specified ranges. Then, for the defined reservoir, the readout ***w***_*out*_ is optimized. Lastly, the trained ESN is evaluated, its performance is compared with other ESN instances and the best performing network is selected. The optimized hyperparameters are described in Table 1.

**Table 1.** Optimized ESN hyperparameters.

| Parameter name | Description | Random search range | Final value (rat \| human) |
| --- | --- | --- | --- |
| Reservoir size | The number of reservoir neurons controls the possible complexity of the performed encoding. Along with the increase in neuron count, the computational costs grow. | [30; 1000] | 343 \| 621 |
| Spectral radius | A key parameter of the network is its spectral radius - the highest absolute eigenvalue of the internal connectivity matrix. The magnitude of this parameter influences the temporal length of the reservoir's memory property<sup>125</sup>. It needs to be set in a way that grants the reservoir the echo state property (ESP), meaning that the ESN's state will become independent of its initial conditions. It has been empirically observed that setting a value lower than 1 usually guarantees that the reservoir will possess the ESP<sup>124</sup>. | [0.6; 1] | 0.74025 \| 0.8741 |
| Connection weights | The choice of a specific spectral radius directly influences the scaling of internal connection weights, however selecting minimal and maximal values for the range from which the weights will be drawn gives control over the ratio and relative strengths of negative and positive links in the network. The same applies to the generation of connections between the input and the reservoir. | Input min:<br>[-1; 0]<br>Input max:<br>[0; 1]<br>Internal min:<br>[-0.5; 0]<br>Internal max:<br>[0; 0.5] | Input min:<br>-0.064538 \| -0.86968<br>Input max:<br>0.37786 \| 0.90886<br>Internal min:<br>-0.15024 \| -0.0045633<br>Internal max:<br>0.20568 \| 0.35022 |
| Network topology | Instead of using a purely random adjacency matrix, a more specific pattern of connections might be preferred. Here, three network topologies were tested:<br>*Small-world* – most links between elements are local, while a small fraction of connections is long-ranged and couples separated clusters of neurons, thus greatly reducing the average path length in the network. The Watts & Strogatz<sup>126</sup> model was used to generate undirected adjacency matrices. These were then converted to directed matrices and the connection weights were drawn from specified ranges. The process was parametrized by:<br>The initial number of every neuron's neighbors. | Topology:<br>[small-world, scale-free, random] | Topology:<br>small-world \| small-world<br>Init. neighbors:<br>56 \| 82<br>Change prob.:<br>0.38912 \| 0.074823 |
|  | ○ Probability of a connection being changed to random (source of long-range connections).<br>○ Probability of keeping a bidirectional connection.<br>*Scale-free* – networks characterized by the power law decay of the probability $P(k)$ that a given node is connected to $k$ other elements. Directed adjacency matrices created using the Barabasi & Albert algorithm<sup>127, 128</sup> have been generated and connection weights have been drawn from specified ranges. Scale-free network parameters:<br>○ The initial number of vertices.<br>○ The number of connections added with every new node.<br>*Random* – no structure is imposed on the connectivity pattern. The only parameter of the adjacency matrix is its connection density. |  | Bidirectional prob.:<br>0.070388 \| 0.54819 |
| Leakage | This parameter controls the fraction of a neuron's activation that will be utilized in the computation of its following state. | [0.01; 1] | 0.31577 \| 0.074823 |
| Time constant | A gain parameter brought about by discretizing continuous time dynamics of leaky-integrator ESN equations<sup>123</sup>. Was fixed to 1 in the human ESN. | [0.1; 1] | 0.080229 \| 1 |
| Bias presence | The input signal might be expanded by an additional bias dimension, providing constant input to all connected units at every time point. | [0; 1] | 1 \| 0 |
| Feedback | The decoded output of the network might be fed back into the network to influence the next state of the reservoir. Feedback presence and the scaling of feedback weights were found using RS. The scaling was drawn from a discrete set of logarithmically spaced values. Initial weights were drawn from the [-1; 1] range. | Presence:<br>[0; 1]<br>Scaling:<br>[10<sup>-14</sup>; 1] | Presence:<br>0 \| 1<br>Scaling:<br>0 \| 10<sup>-10</sup> |
| Washout time | The number of input signals' time points used to drive the reservoir into a dynamical state that is specific to a given input and independent of the initial state (due to the echo state property). The states generated for these points are not used for readout training and prediction. The necessary washout time might be reduced by using an appropriate initial state. | fixed, RS not used | 250 \| 250 |
| Initial state | The initial internal states of all reservoirs' neurons can be set to 0 or to a state resembling the states generated by the data of interest. To generate a state similar to the ones produced by single-vessel signals a part of the training data was fed into the ESN before the training. The mean of all the generated states (distinct for each neuron) was then used as the initial state value. | [0; based on training data] | based on training data \| based on training data |

### GRU and LSTM

The predictions of two other RNN models were compared with ESN’s predictions. Gated recurrent unit (GRU)^82^ and long short-term memory (LSTM)^83, 84^ networks are recurrent neural network architectures designed to tackle the vanishing and exploding gradient problems, which prevented effective learning in networks trained using backpropagation. Both introduce gating mechanisms that control the flow of information into and out of the GRU or LSTM units and allow the networks to capture dependencies at different time scales in the processed data. Like the ESN, both the GRU and LSTM encode each element of the input single-vessel sequence ***x*** into a hidden state vector ***h***(*t*). The GRU computes the following function:

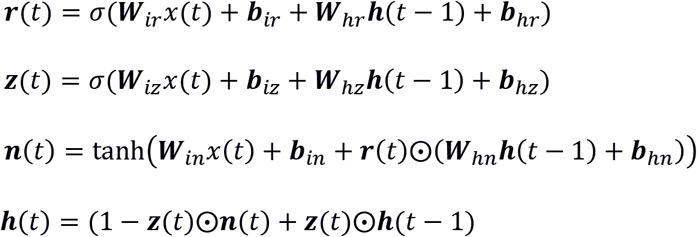

where *σ*(), *tanh* () are the sigmoid and hyperbolic tangent functions, ***r***, ***z***, ***n*** are the reset, update and new gates, ***W****s* are matrices connecting the gates and ๏ is the elementwise product. LSTM’s encoding looks as follows:

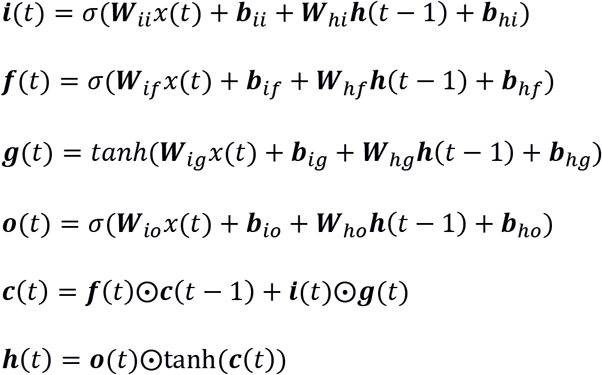

where ***c*** is the cell state, ***i***, ***f***, ***g***, ***o*** are the input, forget, cell and output gates. As in the ESN, in both GRU and LSTM networks, a linear readout was used to generate the prediction based on the state vector:

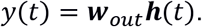

The networks were trained in PyTorch^129^. The same data and cross-validation procedure as during the ESN’s training were used. The hyperparameters were found with Bayesian optimization using the tree of Parzen estimators algorithm (Hyperopt toolbox, n=600)^85, 86^. The optimized hyperparameters have been described in Table 2.

**Table 2.** Optimized backpropagation-RNN hyperparameters.

| Parameter name | Description | Range | Final value<br>(rat \| human) |
| --- | --- | --- | --- |
| Network type | Both network architectures were tested for the rat and human data. | [LSTM, GRU] | LSTM \| GRU |
| Number of layers | Multiple layers of each of the recurrent units could be stacked on top of each other. | [1; 5] | 1 \| 1 |
| Hidden size | Size of the hidden state vector. | [10; 500] | 136 \| 88 |
| Loss function | As CC was the final evaluation metric of networks' performance, it could be used as the cost function instead of the mean squared error (MSE) loss. | [MSE, CC, MSE and CC] | CC \| CC |
| Learning rate | A parameter defining the rate at which network weights were updated during training. | [0.01; 1] | 0.31577 \| 0.074823 |
| L2 | Strength of the L2 weight regularization. | [0; 10] | 0.0025 \| 0.0167 |
| Gradient clipping | Gradient clipping<sup>130</sup> limits the magnitude of the gradient to a specified value. | [yes; no] | yes \| no |
| Dropout | In the case of using a multi-layer RNN, dropout<sup>131</sup> could be set. As in both rat and human datasets 1-layer networks were found, dropout wasn't used. | [0; 0.2] | - \| - |
| Residual connection | Employing a residual connection i.e. feeding the input directly to the linear readout alongside the RNN's hidden state. | [yes; no] | yes \| no |
| Batch size | The number of single-vessel time courses processed by the network in the training stage before each weight update. | [3; 32] | 4 \| 27 |
| Number of epochs | How many times the network processed the whole training dataset during training. | [1; 100] | 67 \| 51 |

### ARMAX

The autoregressive-moving-average model with exogenous inputs (ARMAX)^79^ was used as a comparative prediction method. ARMAX aims to model a time series using autoregressive, moving-average and exogenous input terms. This is depicted in the equation:

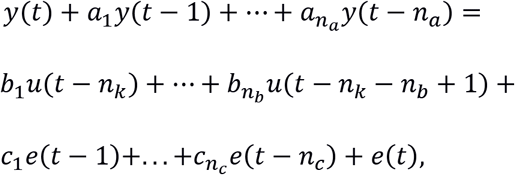

where *y*(*t*) is the model’s output at time *t*; *u*(*t*) is the exogenous input at time *t*; *e*(*t*) is the noise term at time *t*; *n*_*a*_, *n*_*b*_, *n*_*c*_ are the numbers of model’s past outputs, inputs and error terms that influence the current output; *n*_*k*_ is the delay after which the inputs influence the output; *a*_*i*_, *b*_*i*_, *c*_*i*_ are estimated model coefficients. To match the 10 s prediction scheme *n*_*k*_ was set to 10 and the raw inputs and slow oscillation outputs were not shifted. An extensive grid search was performed to find the *n*_*a*_, *n*_*b*_, *n*_*c*_ values that led to the best predictions. All combinations of *n*_*a*_, *n*_*b*_, *n*_*c*_ values ranging from 1 to 50 with a step of 1 and from 1 to 150 with a step of 5 were evaluated to estimate the model’s coefficients *a*_*i*_, *b*_*i*_, *c*_*i*_. Exactly the same data as in ESN’s case were used for training and evaluation and the best set of *n*_*a*_, *n*_*b*_, *n*_*c*_ values was also found through cross-validation. MATLAB *armax* and *forecast* functions were used to find the coefficient values and evaluate the models. ARX and ARIMAX models were also tested but yielded worse performances, hence are not reported.

### Experimental procedures

Single-vessel fMRI data acquired from 6 rats and 6 human subjects have been previously published^24^. The rats were imaged under alpha-chloralose anesthesia. For details related to the experimental procedures refer to^24, 132^.

### Rat MRI data acquisition

The measurements have been performed using a 14.1 T/26 cm horizontal bore magnet (Magnex) interfaced with an Avance III console (Bruker). To acquire the images a 6 mm (diameter) transceiver surface coil was used.

### bSSFP rs-fMRI

The balanced steady-state free precession (bSSFP) sequence was used to acquire 2-5 trials of BOLD rs-fMRI for every rat. Each run had a length of 15 minutes with a one slice repetition time of 1 s. The bSSFP parameters were: TE, 3.9 ms; TR, 7.8 ms; flip angle (FA), 12°; matrix, 96 × 128; FOV, 9.6 × 12.8 mm; slice thickness = 400 µm; in-plane resolution = 100 × 100 µm^2^.

### MGE A-V map acquisition in rats

To detect individual blood vessels a 2D Multiple Gradient-Echo (MGE) sequence was used. The sequence parameters were: TR = 50 ms; TE = 2.5, 5, 7.5, 10, 12.5 and 15 ms; flip angle = 40°; matrix = 192 × 192; in-plane resolution = 50 × 50 µm^2^; slice thickness = 500 µm. The second up to the fifth echoes of the MGE images were averaged to create arteriole-venule (A-V) maps^67^. The A-V maps enable identifying venule voxels as dark dots due to the fast T2* decay and arteriole voxels as bright dots because of the in-flow effect.

### Human MRI data acquisition

Data from six healthy adult subjects (male, n = 3; female, n = 3; age: 20 - 35 years) were acquired using a 3-T Siemens Prisma with a 20-channel receive head coil. BOLD rs-fMRI measurements were performed using an EPI sequence with: TR = 1,000 ms; TE = 29 ms; FA = 60°; matrix = 121 × 119; in-plane resolution = 840 µm × 840 µm; 9 slices with thicknesses of 1.5 mm. Image acquisition was accelerated with parallel imaging (GRAPPA factor: 3) and partial Fourier (6/8). Subjects had their eyes closed during each 15 minute trial. Respiration and pulse oximetry were simultaneously monitored using the Siemens physiologic Monitoring Unit (PMU).

### Data preprocessing

All data preprocessing was done using MATLAB and the Analysis of Functional Neuro Images (AFNI) software package^133^. The functional data were aligned with the A-V map using the mean bSSFP template and the *3dTagAlign* AFNI function with 10 tags located in the venule voxels. Other details of the preprocessing procedure are reported in a previous study^90^. No spatial smoothing was done at any point.

### Localization of individual veins

To localize venule voxels in A-V maps, local statistics analysis and thresholding were performed using AFNI. First, for each voxel, the minimum value in a 1 voxel-wide rectangular neighborhood was found. Then, the resulting image was filtered with a 10 voxel rectangular rank filter and divided by the size of the filter. Finally, the image was thresholded to create a mask with vein locations. For human data, the mean of EPI time series was used instead of the A-V map.

### ICA identification of vascular slow oscillations

To extract signals only from vessels exhibiting strong slow oscillations an additional independent components analysis (ICA)-based mask was combined with the described above vessel localization method. The functional rs-fMRI data were processed using the Group ICA of fMRI Toolbox (GIFT, http://mialab.mrn.org/software/gift) in MATLAB. First, principal component analysis (PCA) was employed to reduce the dimensionality of the data. PCA output was used to find 10 independent components and their spatial maps using Infomax ICA^70^. If a component exhibiting slow oscillations predominantly in individual vessels had been found, it was thresholded and used together with the vascular mask to identify vessels of interest and extract their signals.

### Frequency normalization

To normalize the data, power density estimates of signals’ high-frequency components were used. Every time course had its mean removed and was divided by the mean PSD of its frequency components higher than 0.2 Hz. The 0.2 Hz point was chosen, as above this value spectra of extracted signals were centered on a horizontal, non-decaying line. Performing the division brought the mean PSD of high-frequency components to a common unit baseline for all signals.

This allowed to better compensate for different signal strengths across trials than when scaling the data using minimal and maximal values. Additionally, the relative strength of flatter signals and those exhibiting stronger low-frequency oscillations was better preserved when compared to variance normalization. Ultimately it also improved prediction performance.

### Power spectrum analysis

The spectral analysis was performed in MATLAB. To compute the power spectral density estimates (PSDs) of utilized signals we employed Welch’s method with the following parameters: 1024 discrete Fourier transform points; Hann window of length 128; 50% overlap.

### Filtering

To obtain target signals, single-vessel time courses were bandpass filtered in MATLAB using *butter* and *filtfilt* functions. The frequency bands (0.01-0.1 for human and 0.01-0.05 for rat data) were chosen based on the PSD curves of single-vessel and ICA time courses.

### Surrogate data generation

Surrogate data methods are primarily used to measure the degree of nonlinearity of a time series^74^. They allow creating artificial time courses that preserve basic statistics of original data like the mean, variance and autocorrelation structure. In this study, Fourier based surrogate signals were generated for each single-vessel time course using the iterative amplitude adjusted Fourier transform (IAAFT) algorithm^75^.

To create a surrogate control, a list of a signal’s amplitude-sorted values and the complex magnitudes of its Fourier frequency decomposition need to be saved. First, the original signal is randomly reordered. The complex magnitudes of the shuffled signal are replaced by the stored values of the original signal with the new phases being kept. This changes the amplitude distribution. To compensate for this, the new signal’s sorted values are assigned values from the stored ordered amplitude distribution of the source signal (the new signal is only sorted for the assignment, its order is restored afterwards). In turn, matching the amplitudes modifies the spectrum, so the complex magnitude and amplitude matching steps are repeated and the modified phases of the resulting signal are kept through iterations.

The iteratively generated signals had the same amplitude distribution as the source data and extremely similar amplitudes of the power spectrum. However, the phases of their complex Fourier components were randomized.

### HCP data – preprocessing

Data from 3279 15-minute sessions of rs-fMRI acquired by the Human Connectome Project (HCP)^68^ were used to extract V1 signals and compute whole-brain correlation maps. The data set was preprocessed^134, 135^, had artifacts removed via ICA+FIX^136, 137^ and was registered to a common space^77, 138^ by the HCP. The data were resampled from the original 0.72 s sampling rate to match the 1 s TR of our in-house datasets.

### HCP data – ROI signal extraction

The multi-modal cortical parcellation^77^ was used to extract 180 ROI signals per hemisphere. The DMN ROI was based on the DMN ROI specified in Yeo et al.^139^. Task-positive regions were labeled according to Glasser et al.^77^. Subcortical structures were extracted using the Connectome Workbench^140^.

### HCP data –ICA parcellations

ICA spatial maps and their corresponding time courses for each rs-fMRI session were obtained from the S1200 Extensively Processed fMRI Data released by HCP. The spatial maps are based on group-PCA results generated using MIGP (MELODIC’s Incremental Group-PCA)^141^. Spatial ICA was applied to the group-PCA output using FSL’s MELODIC tool^142, 143^. To derive component-specific time courses for each session, the spatial maps were regressed against the rs-fMRI data^144^. In this work, we used results from the 15-component decomposition. Not all used rs-fMRI sessions had an ICA time course available (initial group sizes in the seed-based analysis were n_top_=202 and n_bottom_=207; for ICA seed-based analysis the sizes were n_top_=195 and n_bottom_=203).

### Cross-correlation

MATLAB *xcorr* and *zscore* functions were used to compute cross-correlation. Lag times were computed between predictions and desired outputs. Positive lags correspond to delayed predictions and negative lags to too early predictions. The input signal has an additional 10 s shift.

### Statistical tests

The statistical significance of the difference between real/surrogate and ESN/ARMAX prediction scores was verified using a paired t-test (MATLAB *ttest* function). To determine differences between seed-based correlation maps and PSDs two-sample t-tests were applied (MATLAB *ttest2* function). The results have been controlled for false discovery rate with adjustment^145, 146^. Fisher’s z-transform has been applied to all correlation values before conducting statistical tests. P values <0.05 were considered statistically significant.

## ACKNOWLEDGMENTS

This research was supported by NIH Brain Initiative funding (RF1NS113278-01), German Research Foundation (DFG) SPP-1655 and Yu215/3-1, BMBF 01GQ1702, internal funding from Max Planck Society. We thank Dr. R. Pohmann and Dr. K. Buckenmaier for technical support; Dr. E. Weiler, Dr. P. Douay, Mrs. R. König, Ms. S. Fischer, and Ms. H. Schulz for animal/lab maintenance and support; the Analysis of Functional NeuroImages (AFNI) team for software support.

## AUTHOR CONTRIBUTIONS

X.Y., F.S. designed the research; X.Y., Y.H., F.S. acquired the data; F. S., X.Y. developed the methods and performed data analysis; T.S. provided conceptual and methodological support, F.S., X.Y., T.S. wrote the paper.

---

**Supplementary fig. 1.**
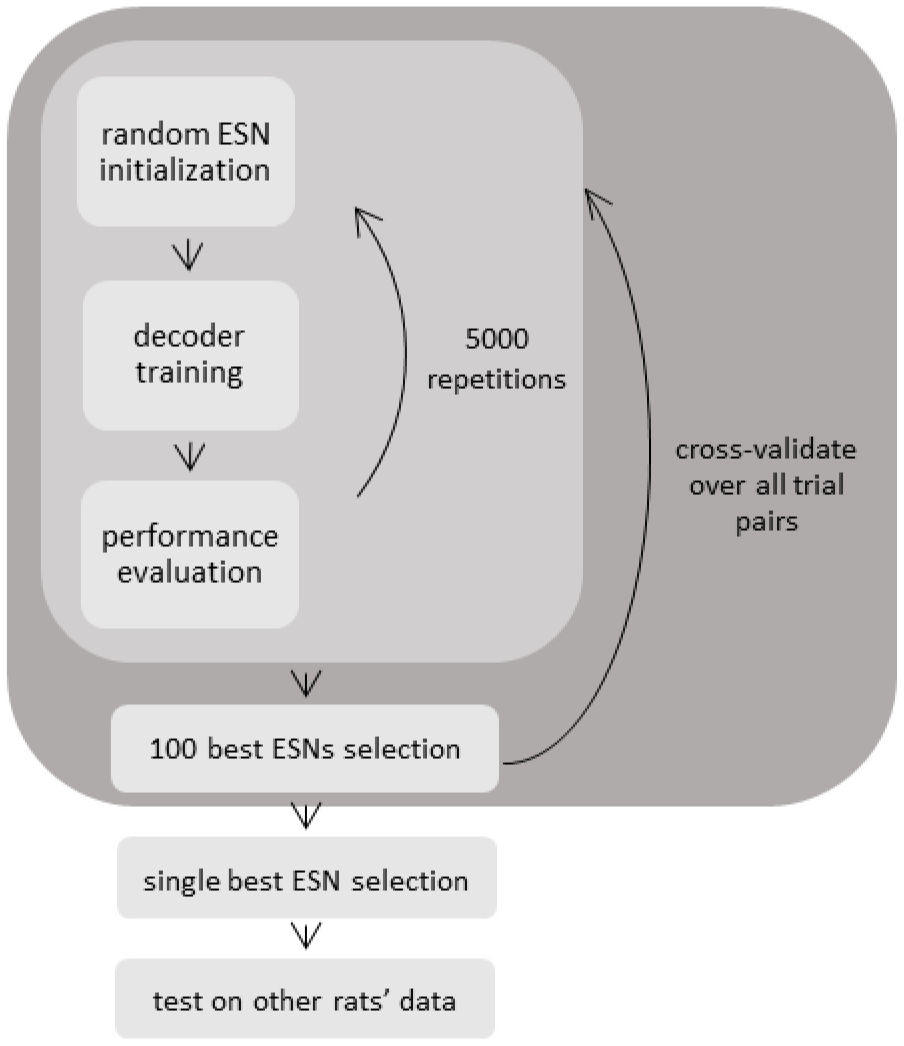
ESN hyper-parameter optimization using random search. For each possible 3+1 trial division (3 training trials + 1 test trial) network parameter values are drawn randomly from pre-specified ranges 5000 times. For each drawn parameter set a reservoir is generated and an ESN is trained and evaluated. For each 3+1 division the 100 best performing ESNs are cross-validated to select a single ESN for predicting other rats’ data.

**Supplementary fig. 2.**
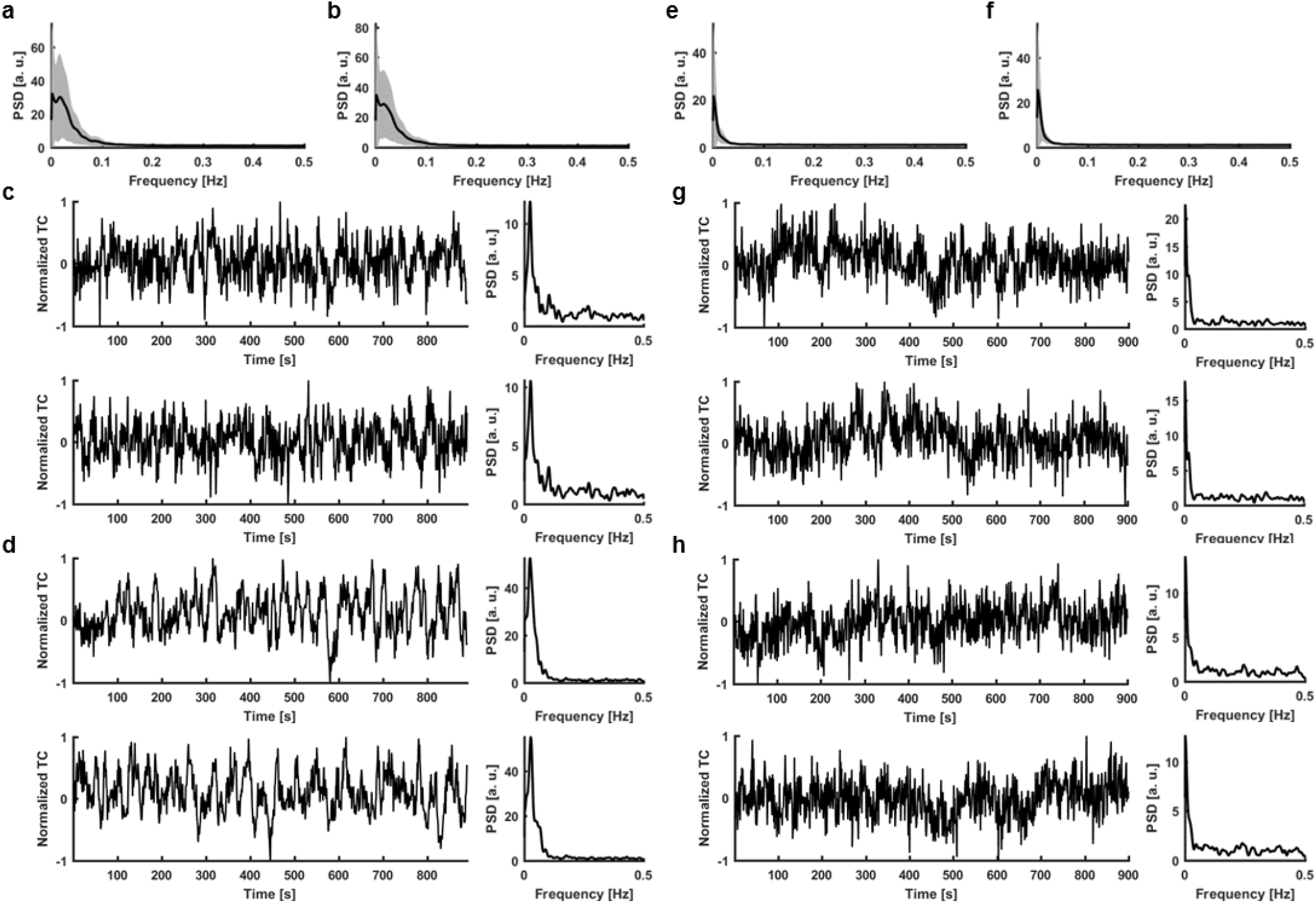
Surrogate data generation. **a**, Mean PSD of all human vessel time courses used in the analysis (shaded area – s.d.). **b**, Mean PSD of all surrogate control time courses generated for comparison with human signals (shaded area – s.d.). **c**, A real human signal and its PSD (top) matched with the generated surrogate control and its PSD (bottom). **d**, Same as c. **e**, Mean PSD of all rat vessel time courses used in the analysis (shaded area – s.d.). **f**, Mean PSD of all surrogate control time courses generated for comparison with rat signals (shaded area – s.d.). **g**, A real rat signal and its PSD (top) matched with the generated surrogate control and its PSD (bottom). **h**, Same as g.

**Supplementary fig. 3.**
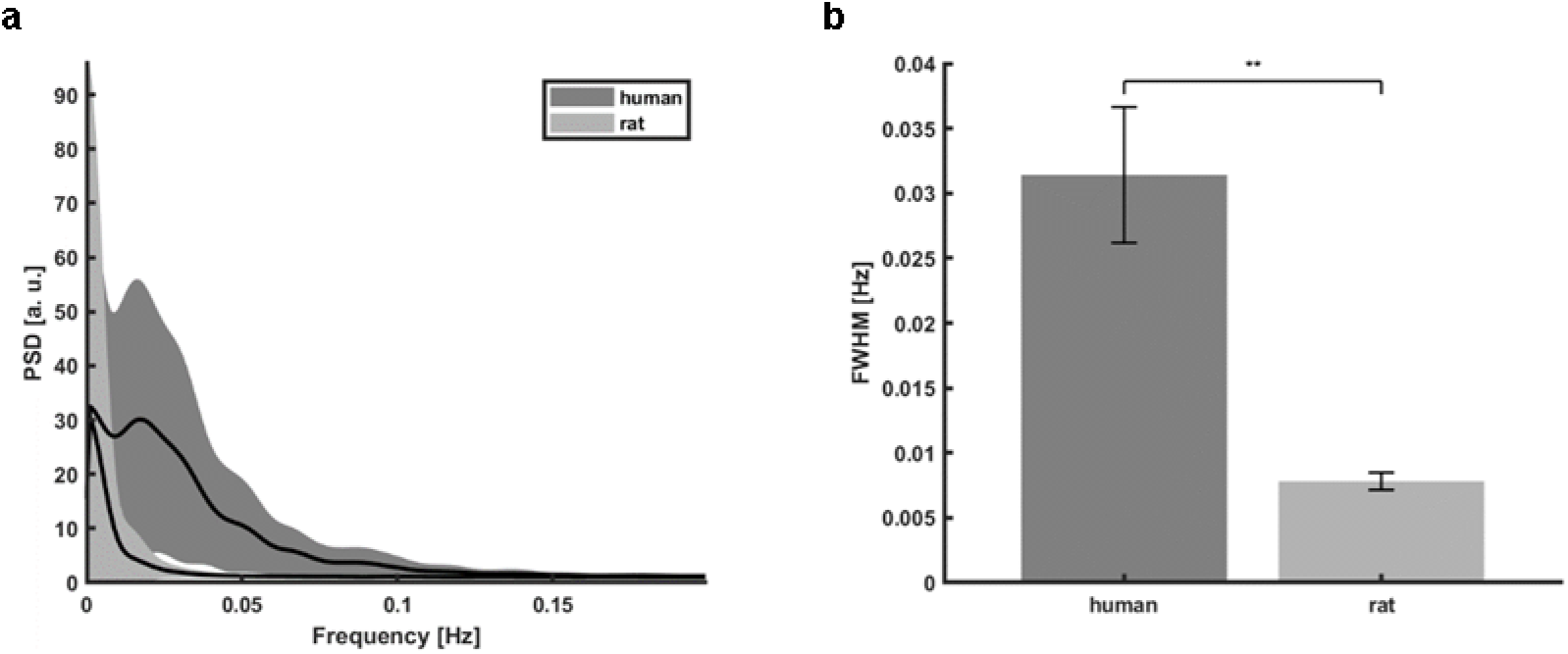
Interspecies PSD difference. **a**, Mean PSDs of all human and rat vessel time courses used in the analysis (shaded areas – s.d.). **b**, Difference of full width at half maximum (FWHM) means of six human subjects’ mean PSDs (0.031 ± 0.01 s.e.m.) and of six rats’ mean PSDs (0.008 ± 0.001 s.e.m.; two sample t-test, p= 0.001).

**Supplementary fig. 4.**
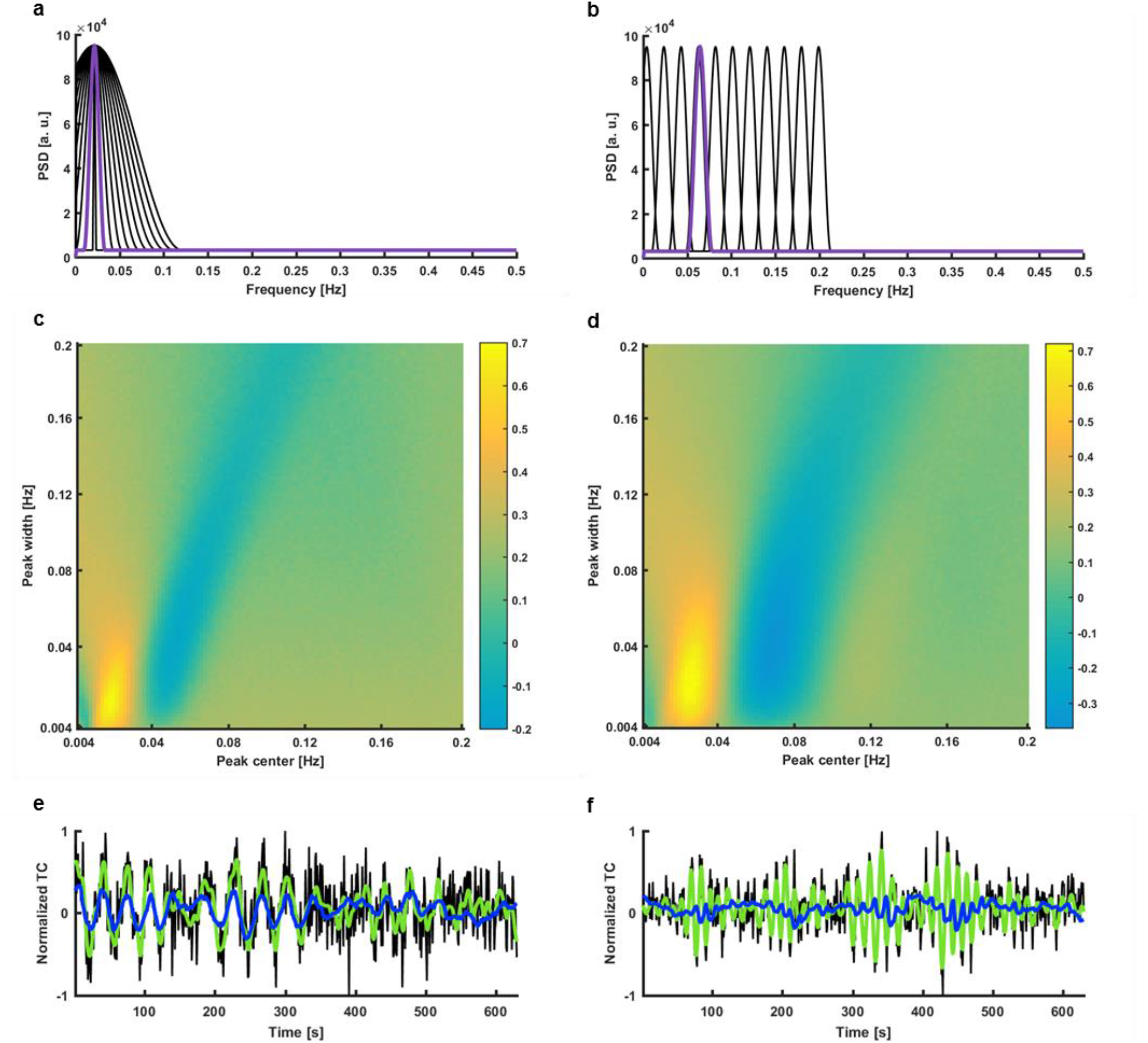
Trained ESN input feature specificity. **ab**, Examples of different artificial PSDs with a fixed peak location and varying peak width (a) or fixed peak width and varying peak location (b) used to generate synthetic time courses with specified spectral features. **cd**, Grid displaying the mean prediction scores of time courses generated for each center location – width pair. Values for both the peak width and location were ranging from 0.005-0.2 Hz and were evenly spaced by 0.002 Hz. For each pair 100 signals were generated. Every point on the grid represents their mean prediction score (c – rat ESN; d – human ESN). **e**, Prediction plot of a signal generated from the width (0.021 Hz) and peak location (0.025 Hz) pair best predicted by the human ESN (CC=0.72, t_lag_=0; black – raw data, green – target prediction, blue – network output). **f**, Prediction plot of a signal generated from the width (0.041 Hz) and peak location (0.068 Hz) pair worst predicted by the human ESN (CC=-0.33, t_lag_=7; black – raw data, green – target prediction, blue – network output).

**Supplementary fig. 5.**
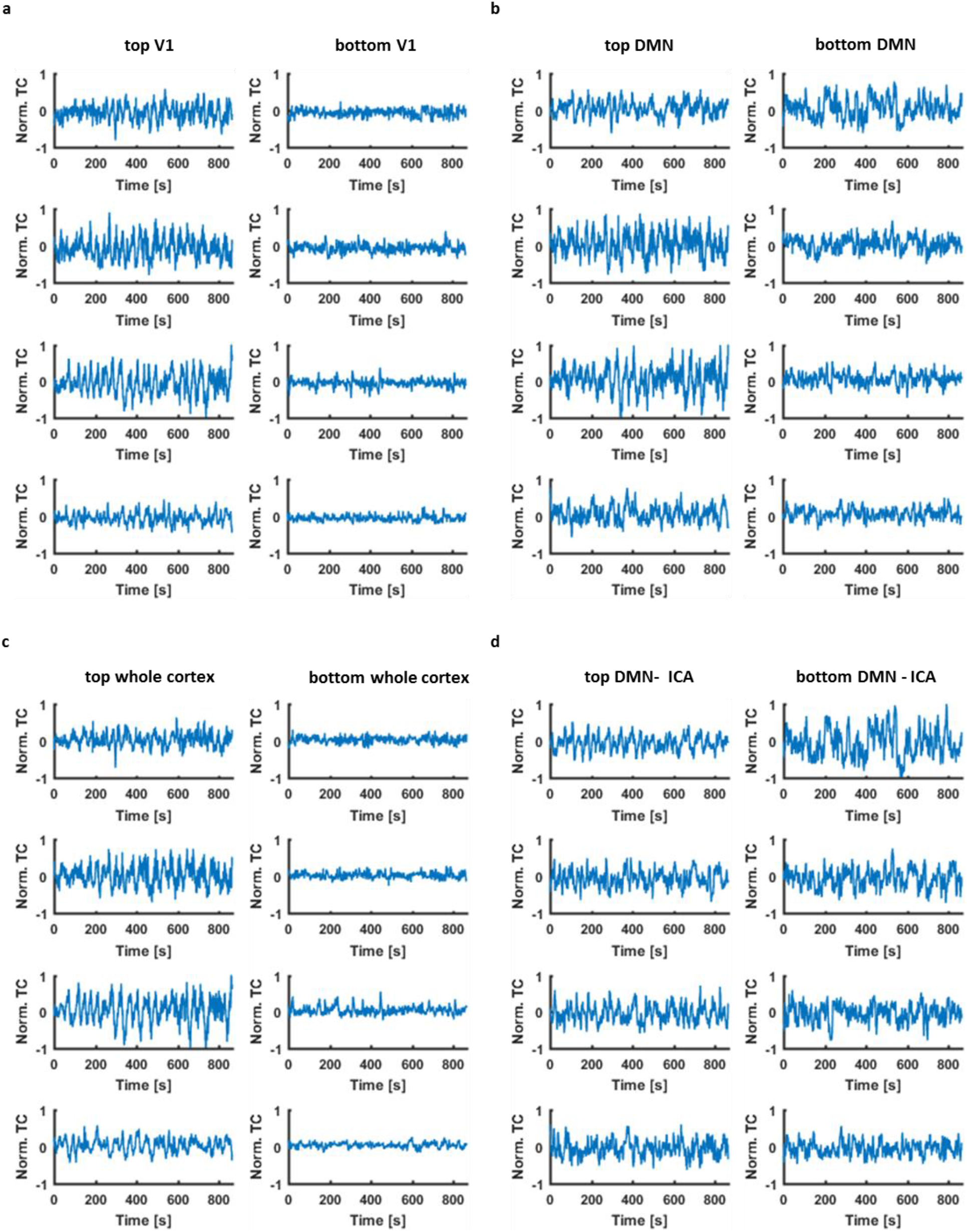
Representative seed ROI signals. Time courses extracted from ROIs used as seeds in the connectivity analysis. Signals from the same ROI have been normalized together. 4 sessions from the “top” and “bottom” groups are shown. **a**, V1 signals. **b**, DMN signals. **c**, Global cortical signals. **d**, DMN ICA signals.

**Supplementary fig. 6.**
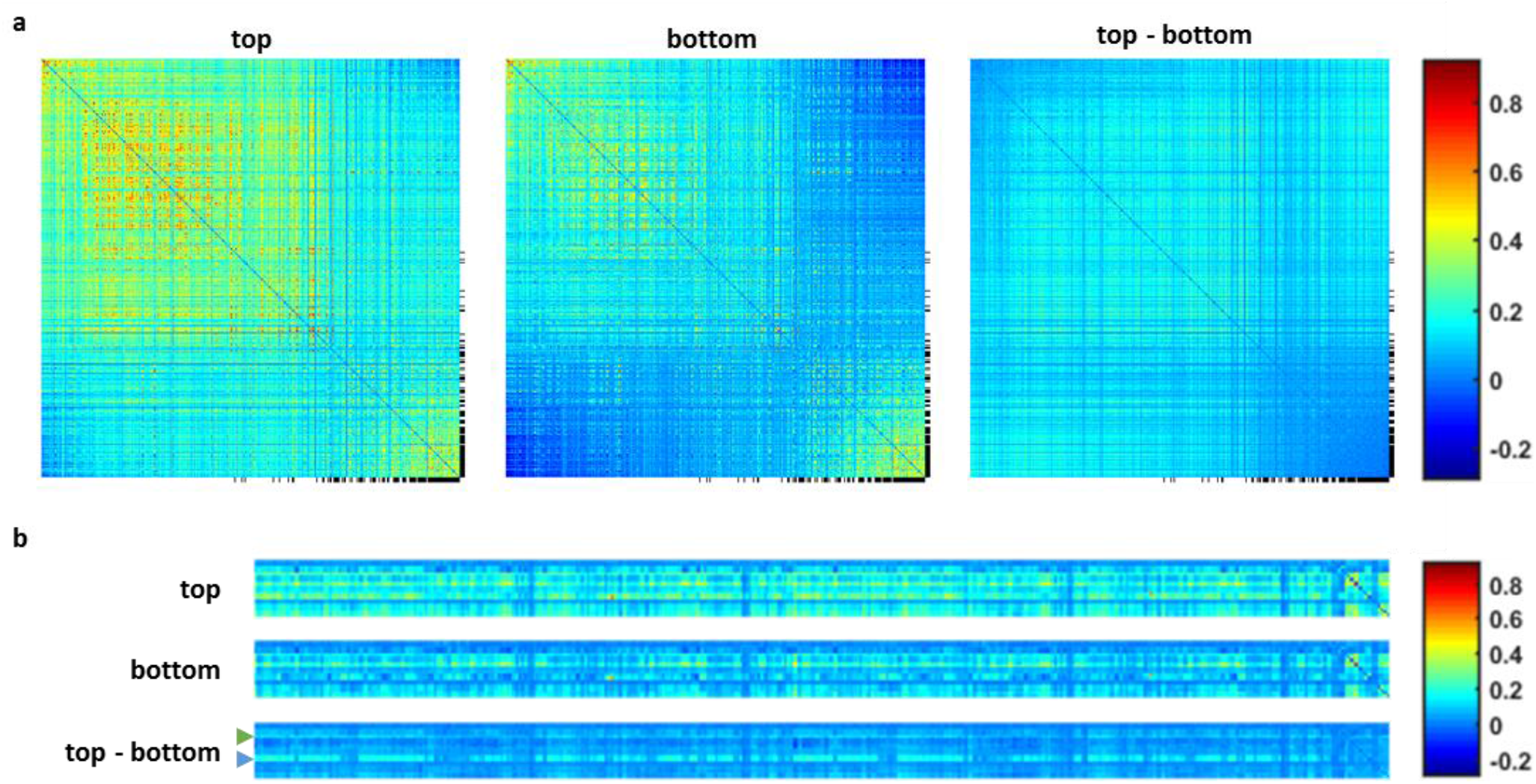
HCP whole brain functional connectivity. **a**, Correlations matrices of 360 cortical ROIs. The matrices have been rearranged based on the order resulting from spectral reordering of the difference matrix. Most of the cortex displays an increased synchrony in the well predicted sessions. Exceptions are the DMN (black markers) ROIs, which despite being more synchronized with the global signal, don’t show an increase in internal connectivity. The values on the diagonal have been set to 0. **b**, Correlation matrices of 19 subcortical ROIs and the cortex. Brainstem (blue arrow) and hippocampus (green arrow) show increased synchrony with the global signal. The rightmost part of the matrices shows internal subcortical connectivity.

**Supplementary fig. 7.**
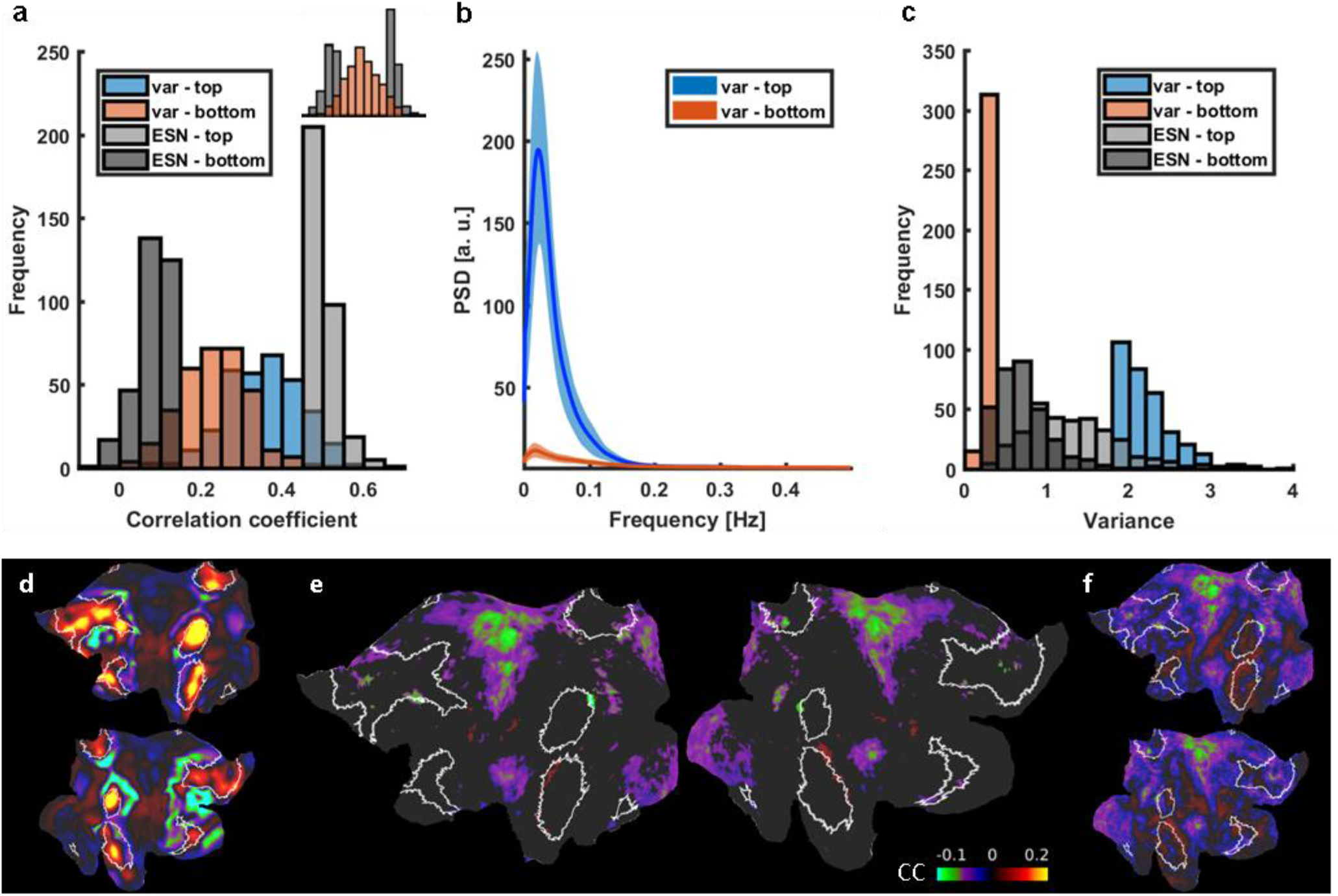
Variance based classification. **a**, Prediction score histograms of the 5% best (light gray) and worst (dark gray) predicted sessions contrasted with prediction scores of signals with top 5% highest (blue) and lowest (red) variance. Variance levels aren’t conclusive of ESNs performance. *Top right:* same data with groups merged. **b**, Mean PSDs of signals with the top 5% highest (blue) and lowest (red) variance. Shaded areas show standard deviations. **c**, Histogram of variance-based (red and blue) and ESN-based (gray) group variance values. Predictions scores of signals having a high or low variance are distributed across the whole range of CC values. **d**, The spatial ICA component (white borders highlighting the DMN), whose time courses have been used to generate the variance-based difference connectivity map. **e**, The variance based differential (“top”-”bottom”) connectivity pattern. It doesn’t resemble the ESN-based DMN internal connectivity reduction. Nodes in which the difference was insignificant are masked. **f**, Same as e but without the mask.

**Supplementary fig. 8.**
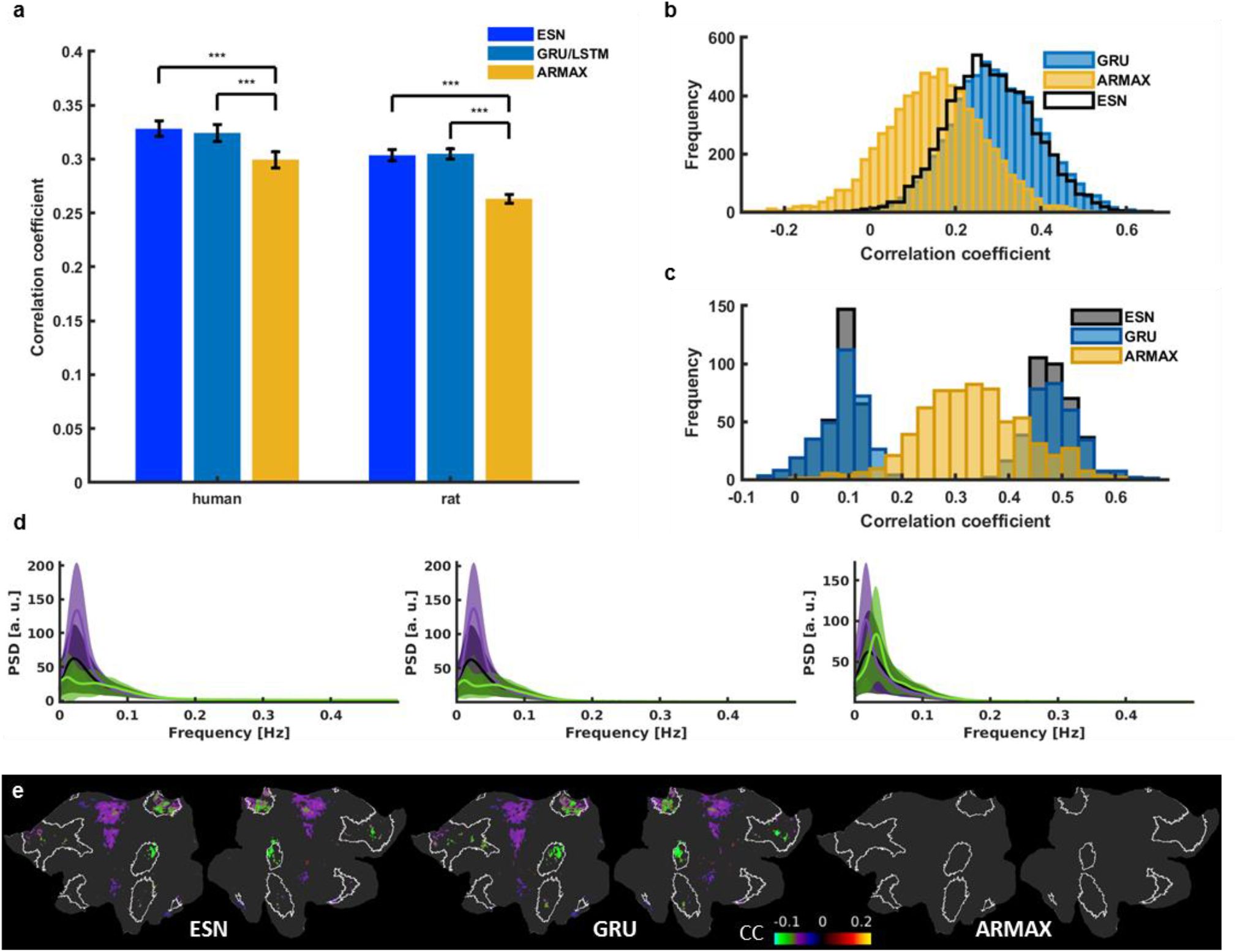
Comparison of different methods’ prediction results. **a**, Mean prediction scores of all in-house human and rat vessel signals obtained using the best ESN, GRU (human data), LSTM (rat data) and ARMAX models. Significantly higher scores (paired-sample t-test, p_H-ESN_=2.84*10^-9^, p_R-ESN_=1.39*10^-23^, p_H-GRU_=8.94*10^-9^, p_R-LSTM_=2.68*10^-27^) obtained by recurrent neural networks than ARMAX in both human (CC_ESN_ = 0.328 ± 0.01; CC_GRU_ = 0.324 ± 0.01; CC_ARMAX_ = 0.299 ± 0.01; mean ± s.e.m.) and rat cases (CC_ESN_ = 0.304 ± 0.01; CC_LSTM_ = 0.305 ± 0.01; CC_ARMAX_ = 0.263 ± 0.01; mean ± s.e.m.). **b**, GRU and ARMAX histograms of prediction scores of 6558 single-hemisphere V1 ROI signals extracted from HCP data. ARMAX predictions are much worse those of the ESN. GRU and ESN prediction score distributions largely overlap. **c**, Histograms showing how much the 5% of best and worst ESN-predicted sessions overlap with the 5% best and worst ARMAX and GRU predictions. ESN and ARMAX predictions show little correspondence. The same sessions were well and poorly predicted by both recurrent networks. **d**, Mean PSDs of time courses whose predictions obtained the bottom 5% (green) and top 5% (violet) scores (left – ESN; middle – GRU; right – ARMAX). Shaded areas show s.d. ARMAX shows less sensitivity to low-frequency oscillatory power compared to the recurrent neural networks. **e**, Flattened cortical maps showing the difference between the mean DMN-ICA-seed-based correlation maps of the “top” and “bottom” groups obtained using the three prediction methods. DMN ROIs are marked by white borders. Nodes in which the difference was insignificant are masked. GRU spatial patterns show significant group differences in the same areas as ESN maps. ARMAX maps don’t show any significant differences. These results suggest that ARMAX prediction accuracy isn’t brain state dependent and that ESN and GRU are tuned to the same features of brain dynamics.

